# The TRPV Channel OSM-9 is Required Non-Cell Autonomously for Sleep-Dependent Olfactory Memory

**DOI:** 10.1101/2020.11.27.400747

**Authors:** Kelli L. Benedetti, Mashel Fatema A. Saifuddin, Julia M. Miller, Rashmi Chandra, Kevin Daigle, Alec Chen, Christine Lin, Angel Garcia, Burhanuddin Calcuttawala, Angelica Tovar, Jackson Borchardt, Kevin Daigle, Raymond L. Dunn, Julia A. Kaye, Saul Kato, Bo Zhang, Maria E. Gallegos, Torsten Wittmann, Noelle D. L’Etoile

## Abstract

Memory, defined as an alteration in behavior towards a stimulus that follows as a consequence of experience, arises when a sensory stimulus is encountered at the same time that the animal experiences a negative or positive internal state. How this coincident detection of external and internal stimuli stably alters responses to the external stimulus is still not fully understood, especially in the context of an intact animal. One barrier to understanding how an intact biological circuit changes is knowing what molecular processes are required to establish and maintain the memory. The optically accessible and compact nervous system of *C. elegans* provides a unique opportunity to examine these processes. *C. elegans* can remember an odor such as butanone when it is paired with a single negative experience and the transient receptor potential (TRP) OSM-9/TRPV5/TRPV6 channel is known to be required for this memory. The multiple gating mechanisms of TRPV channels give them the potential to be the coincidence detectors required to integrate internal state and external stimuli. Here, we report that this TRPV channel is also required for acquisition and possibly consolidation of sleep-dependent, long-term memory of butanone. We find that in the anterior ganglion, endogenous GFP-tagged OSM-9 is expressed in the paired AWA olfactory neurons, the ASH nociceptive neuron pair, the mechanosensory OLQ tetrad, and the paired ADF and ADL sensory neurons. In these cells, OSM-9 protein is concentrated in the sensory endings, dendrites, and cell bodies, but excluded from the neurites in the nerve ring. In the tail, OSM-9 is expressed in the nociceptive phasmid neurons PHA and PHB, possibly PQR as well as PVP. In the midbody, it is possibly expressed in the mechanosensitive PVD neuron. It is notably absent from the AWC pair that are required for butanone attraction. Chronic loss of OSM-9 in a subset of ciliated neurons that do not include AWA interferes with consolidation but not learning. Because OSM-9 is expressed and required in sensory neurons that are not needed for butanone chemosensory behavior, two interpretations are possible. The first, is that OSM-9 loss leads to gain of function or neomorphic behavior of these cells that are extrinsic to the primary sensory circuit and their new activity interferes with acquisition and consolidation of memory. The second is that loss of OSM-9 leads to a loss of function phenotype in which the wild type function of these cells is diminished and this function is required for memory consolidation.

**Author summary:** How organisms learn from their environment and keep these memories for the long term ensures their survival. There is much known about the regions of the brain and the various proteins that are essential for memory, yet the exact molecular mechanisms and dynamics required are not known. We aimed to understand the genetics that underlie memory formation. We tested a gene that encodes a transient potential receptor channel vanilloid channel, which is similar to the channels we have that sense spicy foods and other harmful cues. Our studies have shown that this gene is required for the animal to be able to acquire and perhaps consolidate olfactory memory. This protein is not expressed in the sensory neurons that respond to the odor that is memorized or in other downstream interneurons in the odor-sensation circuit, but it is expressed in a distinct set of sensory neurons. This indicates that long-term memory involves wild type behavior of a wider array of sensory neurons than is required for the primary sensation. These channels are also implicated in neurological disorders where memory is affected, including Alzheimer’s disease. Understanding how memory formation is affected by cells outside the memory circuit might provide testable hypothesis about what goes awry in Alzheimer’s disease.

## Introduction

Learning and memory are vital to survival. Organisms must update their knowledge of their surroundings based on environmental cues. One way to promote learning and memory is through neural plasticity, which involves changes at the structural, molecular, and functional levels. Pioneering studies on the *Aplysia* gill-withdrawal reflex demonstrated the importance of synaptic plasticity on learning and memory through *in vitro* pharmacology and physiology experiments on dissected circuits [1]. Learning and memory are thought to be driven by two types of synaptic plasticity: the first is Hebbian, which includes synaptic long-term potentiation (LTP) and long-term depression (LTD) [2], and the second is non-Hebbian homeostatic compensation, which allows a neuron to return to a set point [2, 3]. A large number of molecules implicated in these processes include factors that reorganize the actin cytoskeleton, affect trafficking, insertion and localization of receptors and cell adhesion molecules, localization of newly translated products, kinase function, and transcription of new genes [4]. As deep as our understanding of these synaptic processes may be, we are still missing the roles of some molecular players for memory in an intact organism. One such player is the TRP channel.

Transient receptor potential (TRP) ion channels were first discovered in the *Drosophila* eye photoreceptor cells [5–7], but have since been found in mammals and even in the single-celled yeast. Classically, TRP channels have been well established as magnesium-, sodium-, and calcium-permeable cation channels that can be activated by and simultaneously integrate multiple intracellular or extracellular stimuli to drive downstream signaling and membrane depolarization. These stimuli include, but are not limited to, pain, temperature, pH, injury, osmolarity, and cytokines [8]. There are seven TRP channel families and all share a similar topology: six transmembrane helices, a short pore helix, and a cation-permeable pore loop region. The TRP families differ by the number of their N-terminal ankyrin repeat domains and whether they have a C-terminal TRP box helix, which is a long helix parallel to the membrane [9]. The TRPV family members are generally considered to be first-line nociceptive sensory receptors and include TRPV1-6. TRPV1 is stimulated by vanillin, vanillic, endogenous lipids like the *N-*acylamides (e.g. endocannabinoids such as anandamide), high temperature (≥43°C), pain, ethanol, low pH, black pepper, garlic (allicin), cannabidiol (CBD), arachidonic acid, resiniferatoxin, and spicy foods, driven by the agonist capsaicin [8–11]. The six ankyrin repeats in TRPV1 each contain two short anti-parallel helices and a finger loop and have been shown to interact with the proteins calmodulin and PI3K [9, 12]. Upon activation, the TRPV1 channel allows calcium ions to enter the cell at the plasma membrane through the pore loop region, ultimately leading to desensitization of the channel and a subsequent refractory period [9]. Although the sensory role of TRPV proteins is well established, other roles of the TRPV channels, including the promotion of neural plasticity required for memory formation, remain obscure.

Within the central nervous system, TRP channels have been shown to promote plasticity [14]. For example, TRPV channels are expressed in the hippocampus where they were implicated in slow synaptic transmission [15], LTP[16], and LTD [17]. TRP channels have recently been examined for their role in the synaptic plasticity that underlies memory. For example, the TRPA channel (painless) plays a role in fly courtship associative long-term memory [18]. TRPCs are the canonical TRP channels most related to *Drosophila* TRP channels and are activated by phospholipase C, diacylglycerol, or conformational coupling [19]. In the mammalian brain, the TRPCs function in the hippocampus to drive synaptic transmission and plasticity, and subsequent spatial working memory [20, 21]. In addition to the hippocampus, expression of TRPCs has been uncovered in the temporal lobe, suggesting a role in neural plasticity, learning, and memory. The TRPMs (melastatin), which were originally discovered in melanomas, are widely expressed in mammalian tissues and can be found in intracellular membranes and at the plasma membrane [8]. TRPM4 has also been found to be expressed in the mammalian hippocampus and is important for plasticity and spatial working and reference memory [22]. Several studies have implicated mammalian TRPV1 in memory formation, including fear memory consolidation behavior in mice [16,23,24]. TRPV1 agonists were shown to overcome the ability of acute stress to block spatial memory retrieval in young rats. Likewise, it was shown to overcome the LTP-blocking and LDT-enhancing effects of stress in CA1 slices of the hippocampus [24]. Further dissection of the exact cellular or molecular processes that depend on TRPV1 in order to promote or permit memory could help provide a mechanistic understanding of how learning and memory are driven.

An outstanding question is at which stage of memory formation does TRP channel activity promote or permit memory in its acquisition, consolidation, or recall? In order to study the required genes in a live animal model as it is behaving, unbiased screens for genes required for learning and memory are needed.

Using *C. elegans*, several unexpected genes were identified in screens for short-term circuit plasticity, one encodes the transient receptor potential vanilloid 5 and 6 (TRPV5 and TRPV6) channel, OSM-9 [25]. The *C. elegans* nematode exhibits olfactory classical conditioning: prolonged exposure to innately attractive food-related odors in the absence of food elicits a downregulated response to the odor [25]. The *osm-9* gene was found to be required for short-term olfactory memory in this paradigm. However, the mechanisms and dynamics of TRPV function in long-term memory formation [26, 27] remained unknown. In addition, a role for *osm-9* in long-term sleep-dependent memory had not been not been reported. Examination of the mechanisms and dynamics underlying TRPV-mediated memory in live *C. elegans* would help us understand how these channels act to integrate or permit integration of signaling over time and collaborate with or permit sleep to promote memory formation in a behaving animal.

*C. elegans* provides a good model system to approach this problem. It is genetically-tractable and its nervous system comprises just 302 neurons connected via 7,000 synapses and was the first organism where the entire connectome has been mapped [28, 29]. By contrast, the mammalian nervous system has 100 billion neurons connected by trillions of synapses and the entire neural connectivity map remains unknown [30]. Additionally, the nematode can form long-term associative memories of olfactory stimuli [26]. Furthermore, the neural circuit underlying olfaction is well established [28,31,32]. Importantly, it is transparent which makes possible single cell resolved calcium activity imaging in the live animal. Thus, studying plasticity in the olfactory circuit in *C. elegans* has the potential to provide insight into the mechanisms by which experience changes complex neural circuits to promote memory formation.

The *osm-9* gene is an ortholog of human TRPV5 and TRPV6. TRPV5 and TRPV6, unlike the rest of the TRPV family, have no known ligands, nor are they thermosensitive [33, 34]. They are highly-calcium selective [35] and TRPV5 and TRPV6 are localized to epithelial cells in the kidney and intestine [36] where they are constitutively open at physiological membrane potentials and are modulated by calmodulin in a calcium-dependent manner [35].

Initially, *osm-9*/TRPV5/TRPV6 was discovered in a screen for *C. elegans* mutants that were defective for sensing high osmolarity. It was subsequently studied for its role in diacetyl chemotaxis as well as short-term associative olfactory memory [25, 37]. Later studies revealed that *osm-9* also has a role in sensing noxious stimuli, allowing for SDS avoidance behavior through its expression in the ASH sensory neurons [38]. This avoidance behavior can be phenocopied through heterologous expression of mammalian TRPV1 in ASH and application of the capsaicin agonist [38], where calcium is likely driven into the cell through the pore loop region to elicit depolarization [39]. Polyunsaturated fatty acids, including the TRPV1 endogenous activator arachidonic acid, function upstream of *osm-9* to mediate AWA-dependent olfactory transduction [40]. The endogenous agonist nicotinamide was discovered to activate the heteromeric OSM-9/OCR-4 channel expressed heterologously in *Xenopus* oocytes, and nicotinamide-induced toxicity acts, in the worm, by opening the OCR- 4/OSM-9 channel [41].

The *osm-9* gene has also been shown to play a role in developmental programming and is downregulated transcriptionally after environmental stress and this downregulation causes animals to avoide specific pheromone cues [42, 43]. These changes in *osm-9* expression were shown to be mediated by DAF-3, chromatin remodeling and endogenous siRNA pathways [43]. Thus, *osm-9* has roles in sensory transduction in response to noxious stimuli, can be activated by polyunsaturated fatty acids, and plays a role in learning and memory, just like TRPV1. OSM-9 shares about ∼25-26% protein sequence similarity to human TRPV1, TRPV5, and TRPV6, and also has intracellular N and C termini, six N-terminal ankyrin domains, and six transmembrane domains, with a calcium-permeable pore loop between the fifth and sixth. Thus, studies on *osm-9* may help to uncover conserved mechanisms of learning and memory that are conserved across phyla.

In our classical conditioning paradigm, we focus on the *C. elegans* response to the food-related odorant butanone, which is specifically sensed by the AWC^ON^ olfactory sensory neurons (OSNs) [44, 45]. The *C. elegans* nematode exhibits olfactory classical conditioning: one 80-minute exposure cycle to butanone in the absence of food causes the nematode to downregulate its chemotactic response to that odor. The second messenger for odor sensation within the AWCs is cGMP [46, 47]. Prolonged odor stimulation leads to AIA interneuron circuit activity and PI3K signaling to the AWC neurons to drive accumulation of the cGMP-dependent protein kinase EGL-4 [48] within the AWC nucleus. These events promote memory of the odor being profitless after 80 minutes of conditioning [31,49,50]. Loss-of-function mutations in *osm-9* cause defective learning that AWC-sensed odors are profitless, and defective chemo-, osmo-, and thermosensation promoted by other sensory neurons [37]. Importantly, loss of *osm-9* restores function to mutants that ignore AWC-sensed odors as a consequence of expressing nuclear EGL-4. After EGL-4 nuclear translocation, nuclear endogenous RNAi promotes a dampened response to butanone by transcriptionally-downregulating the *odr-1* gene through heterochromatin remodeling via *hpl-2/HP1*.

We showed that the *osm-9* gene is epistatic to gain-of-function *hpl-2* or *egl-4* [49, 51]. Thus, the TRPV channel *osm-9* is required downstream of both of these nuclear events to block chemotaxis and is thus the most downstream component of this paradigm currently known. However, the role of *osm-9* in this plasticity paradigm is still not understood. We were also curious if *osm-9* plays a role in long-term olfactory memory since its role in short-term olfactory memory is established. We wanted to identify the time during which the *osm-9* gene is required for long-term memory formation, which of its structural components allow this plasticity, what cells it is expressed in and which cells it is required to be expressed in to allow long-term memories to be consolidated.

We showed previously that the 80-minute butanone conditioned response is temporary and is lost after only 30 minutes recovery on food [27]. However, after odor spaced-training, *C. elegans* can exhibit long-term aversive memory phenotypes [27, 52]. Worms subjected to three cycles of 80-minute exposure to the unprofitable odor in the absence of food with recovery periods on food in between each cycle will keep the memory and continue to ignore the odor for much longer periods of time if they are allowed to sleep (at least 24 hours after training) [27].

Here, we find that *osm-9* null mutants can be conditioned to ignore butanone after undergoing a spaced-training protocol. They do not, however, become repulsed from butanone, which is what happens to wild type animals after such training. Long-term memory consolidation also requires *osm-9.* Specifically, we find that *osm-9* mutants lose any initial odor memory they have after only 30 minutes, and it remains lost. The wild-type animals also lose the memory after 30 minutes, but regain it and keep it for at least 16 hours. Memory fragility in the period after conditioning is a hallmark of consolidation [53, 54]. Thus, *osm-9* likely either plays a role in long-term memory consolidation or it is required to prevent interference with this process.

We also find that the C-terminus of OSM-9 is not required for long-term memory acquisition, since removing most of the C-terminal tail does not affect memory. By tagging OSM-9 with GFP at its endogenous locus via CRISPR-mediated editing, we were able to visualize native levels of GFP-OSM-9. GFP-OSM-9 is expressed in eight head neurons in this strain (AWA, ADF, ADL ASH and the OLQs), but none of these are the AWC neuron. This was not expected as AWC is required for butanone chemoattraction [55] and other promoter expression lines had shown expression in this neuron [37].

GFP-OSM-9 is excluded from the nucleus and found concentrated in the sensory endings of AWA and the OLQ neurons. It is found in the dendrites but is absent from the axo-dendritic neurites of the anterior nerve ring. Though TRP channels have been extensively studied as sensory receptors, they have not been well studied as integrators of prolonged stimulation. This work may indicate that OSM-9 and the cells it is expressed in either contribute to memory acquisition and consolidation by acting in sensory neurons that are not required for the naive butanone response or the wild type activity of the cells that express OSM-9 is altered by loss of OSM-9 such that they interfere with memory consolidation.

## Results

### OSM-9 is required for short-term memory after one cycle and multiple cycle bouts of experience

Although it has been previously shown that *osm-9* is required for olfactory short-term memory of food-related odorants (27and Fig 1, S1 Fig), its role in long-term memory formation has not been reported. We utilized a long-term memory spaced-training paradigm in which we performed odor conditioning in three spaced intervals with recovery periods on food in between (S1 Fig). We tested *osm-9(ky10)* null mutants versus wild-type animals and found that *osm-9(ky10)* animals show incomplete short-term odor memory after one cycle of conditioning [25] (Fig 1) and they acquire less memory immediately after spaced-training than wild types (Fig 1) (Figs 1A and 1B).

**Fig 1.**
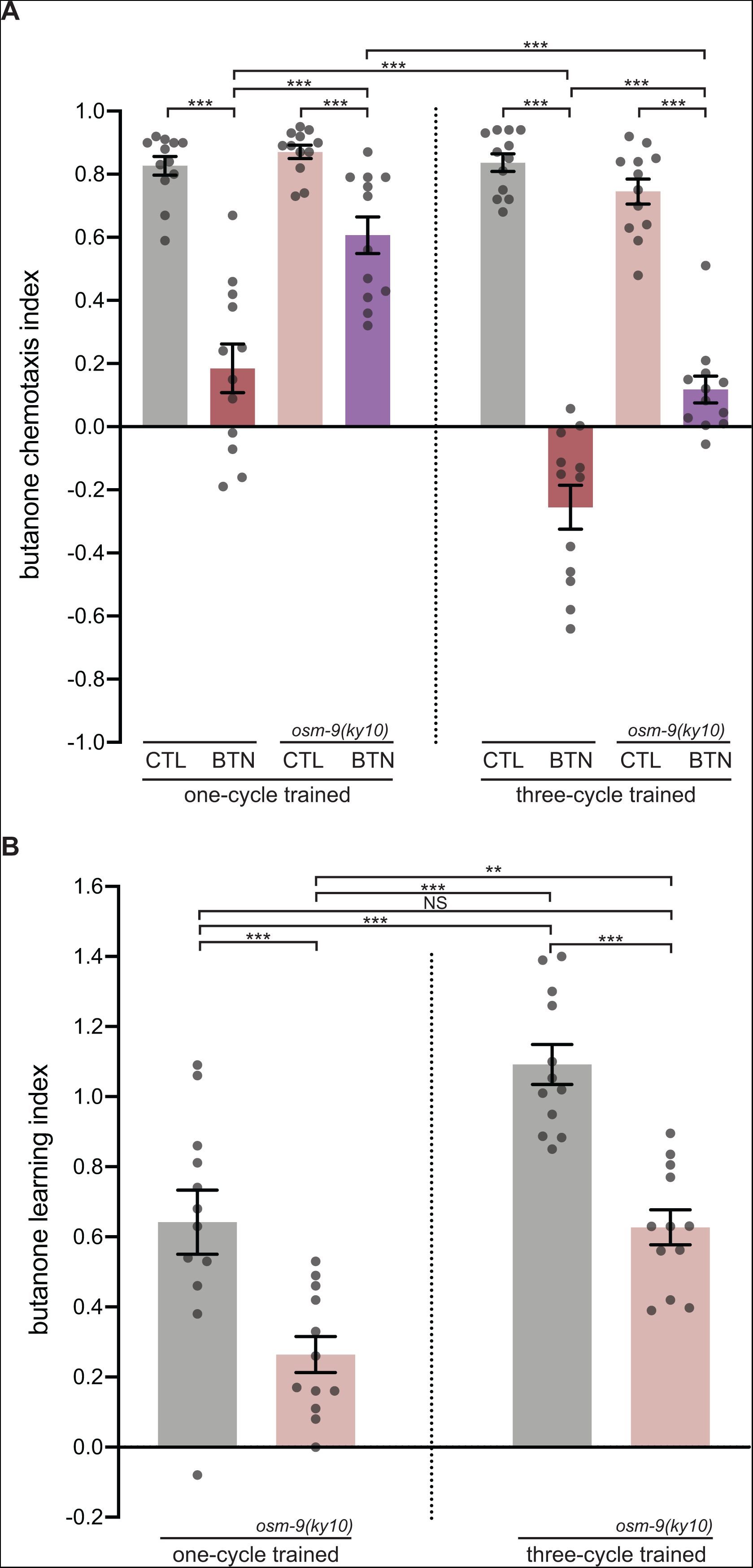
*osm-9* is required for short-term memory, but not memory acquisition post spaced training. (A) Chemotaxis indices for one-cycle odor conditioned and three-cycle odor wild-type vs *osm-9(ky10)* spaced-trained animals. The chemotaxis index (CI) is calculated as the number of worms at the diluent (200-proof ethanol) subtracted from the number at the butanone point source, divided by the total number or worms (excluding any at the origin). Wild-type one- or three-cycle buffer trained (CTL, grey bar), wild-type one- or three-cycle butanone-trained (BTN, red bar), *osm-9(ky10)* one- or three-cycle buffer-trained (CTL, pink bar) and *osm-9(ky10)* one- or three-cycle butanone-trained (BTN, purple bar) are shown, and are denoted this way throughout the rest of the manuscript. “Three-cycle trained” label means 0’ recovery (fifth to eighth bars) and will be annotated this way throughout the paper. N = number of trials where all trials are done on independent days and each grey dot represents an individual assay day, with 50-200 animals/day. The Shapiro-Wilk normality test was performed to determine data distribution, and if data were non-normally distributed, the analysis was done with the Kruskal-Wallis test, an analysis of variance of multiple comparisons for non-parametric data. If pairwise comparisons were normally distributed, then p-values were generated with a Student’s unpaired t-test. If any of the datasets were non-parametric, then p-values were generated with the Mann-Whitney u-test. Hochberg adjustment for multiple comparisons was performed on all p-values generated from data included in the same graph to control Type I statistical error. If the data were normally distributed, then one-way ANOVA was performed, followed by Bonferroni correction of pairwise comparisons. Error bars represent S.E.M. Statistical significance was reported as ***p<0.001, ** p< 0.01, *p<0.05, and NS is p>0.05. Behavioral data throughout the paper are represented in this same way with the same numbers of animals on independent days. Additionally, throughout the paper, the same statistical analysis was performed on the data. Note that for every figure in this manuscript, that the Kruskal-Wallis or one-way ANOVA test was performed and yielded ***p<0.001, unless otherwise noted. The Kruskal-Wallis test was performed. The u-test was performed for one-cycle CTL vs BTN, one-cycle *osm-9(ky10)* CTL vs *osm-9(ky10)* BTN, three-cycle trained *osm- 9(ky10)* CTL vs *osm-9(ky10)* BTN, one-cycle *osm-9(ky10)* BTN vs three-cycle *osm- 9(ky10)* BTN, and three-cycle BTN vs *osm-9(ky10)* BTN. The t-test was performed for one-cycle BTN vs *osm-9(ky10)* BTN, three-cycle CTL vs BTN, and one-cycle BTN vs three-cycle BTN. (B) Learning indices for one-cycle conditioned and three-cycle spaced-trained wild-type vs *osm-9(ky10)* animals. The learning index (LI) is calculated as the chemotaxis index of the BTN animals subtracted from the chemotaxis index of the CTL animals. The higher the learning index, the more the animals have learned and thus kept the BTN memory. Wild type is shown with grey bars and *osm-9(ky10)* with pink bars, and will be denoted this way throughout the manuscript. One-way ANOVA followed by Bonferroni correction was performed for one-cycle wild type vs *osm-9(ky10)*, one-cycle vs three-cycle wild type, one-cycle wild type vs three-cycle *osm-9(ky10),* one-cycle *osm-9(ky10)* vs three-cycle wild type, one-cycle vs three-cycle *osm-9(ky10)*, and three-cycle wild type vs *osm-9(ky10)*.

### OSM-9 is required for consolidation of memory

Wild-type animals lose the memory after 30 minutes of recovery on food, but regain it after two hours of recovery (Fig 2) [27], and keep it for at least 16 hours post training (Fig 3). By contrast, *osm-9* animals lose the memory after only 30 minutes (Fig 2) of recovery on food and never gain it back, as shown by the chemotaxis indices at two and 16 hours of recovery after training (Figs 2 and 3, respectively). This could indicate that *osm-9* is responsible for both acquisition and memory consolidation. Also interesting to note is that although *osm-9(ky10)* animals are able to acquire memory after three cycles of odor training, their chemotaxis behavior looks distinct from wild-type animals – *osm-9(ky10)* animals have an average chemotaxis index closer to zero (0.1, Fig 1A), which looks like wild-type animals after one-cycle of odor training (Fig 1A). By contrast, wild-type animals show an aversion to butanone after three cycles of odor training, with an average chemotaxis index of -0.3 (Fig 1A). This may indicate that *osm- 9* is required for the repulsion induced after odor training. Although repeated spaced odor training can induce memory acquisition in *osm-9(ky10)*, their behavioral state may be different from that observed in wild-type animals. This long-lasting memory has been shown to require sleep [27].

**Fig 2.**
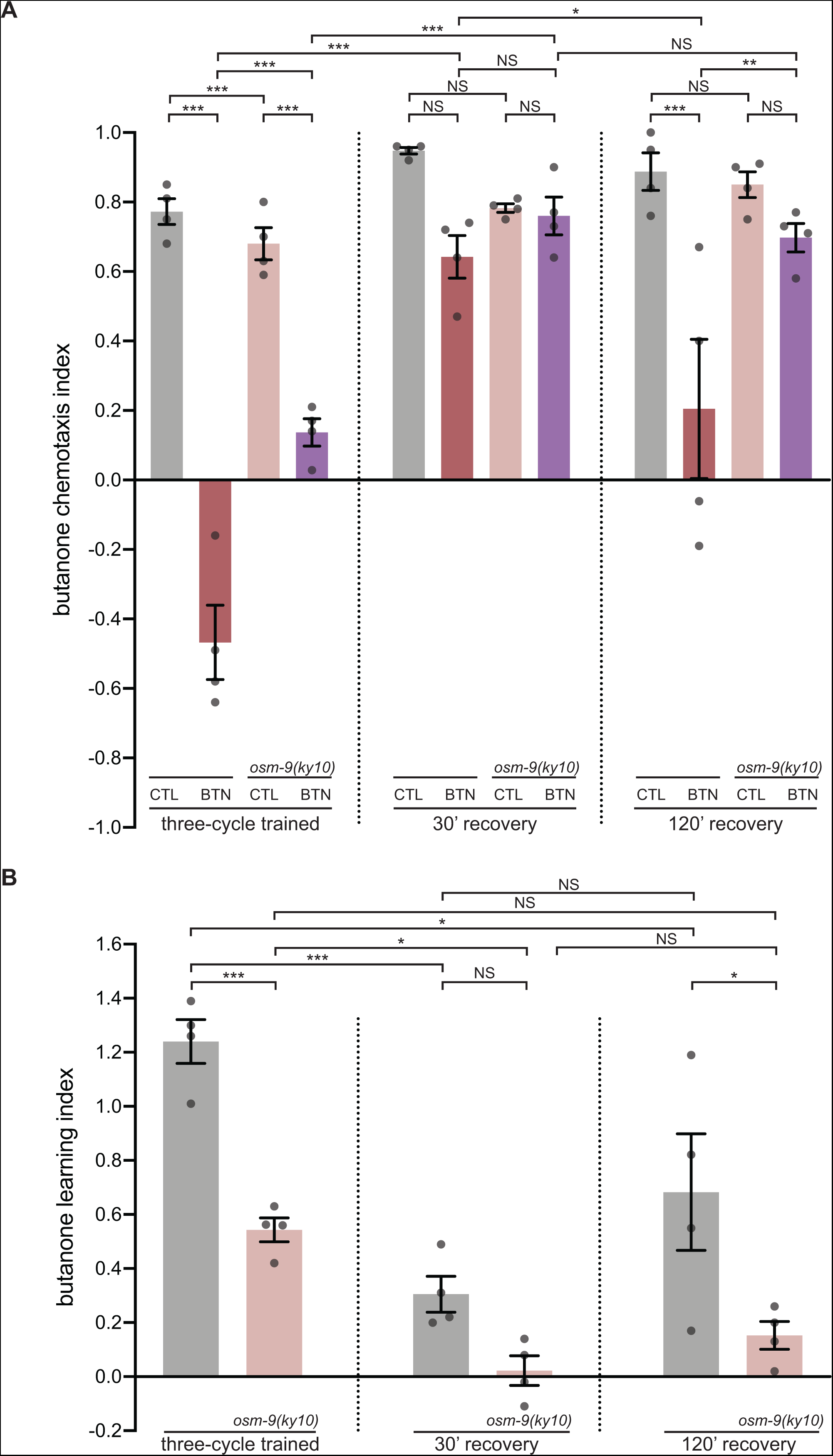
*osm-9* is required for long-term olfactory memory consolidation post spaced training. (A) Chemotaxis indices for three-cycle spaced-trained animals after 0’, 30’ or 120’ post recovery on plates seeded with OP50 as a food source. One-way ANOVA followed by Bonferroni correction was performed for CTL vs BTN, *osm-9(ky10)* CTL vs *osm-9(ky10)* BTN, CTL 30’ vs BTN 30’, *osm-9(ky10)* CTL 30’ vs *osm-9(ky10)* BTN 30’, CTL 120’ vs BTN 120’, *osm-9(ky10)* CTL 120’ vs *osm-9(ky10)* BTN 120’, CTL vs *osm-9(ky10)* CTL, BTN vs *osm-9(ky10)* BTN, CTL 30’ vs *osm-9(ky10)* CTL 30’, BTN 30’ vs *osm-9(ky10)* BTN 30’, CTL 120’ vs *osm-9(ky10)* CTL 120’, BTN 120’ vs *osm-9(ky10)* BTN 120’, BTN vs BTN 30’, *osm-9(ky10)* BTN vs *osm-9(ky10)* BTN 30’, BTN 30’ vs BTN 120’, and *osm-9(ky10)* BTN 30’ vs *osm-9(ky10)* BTN 120’. (B) Learning indices for three-cycle spaced-trained animals after 0’, 30’, or 120’ recovery on food. One-way ANOVA followed by Bonferroni correction was performed for wild-type vs *osm-9(ky10)* after 0’ recovery (compare first two bars), 30’ recovery (compare third and fourth bars), 120’ recovery (compare fifth and sixth bars), wild-type 0’ vs 30’ *osm-9(ky10)* 0’ vs 30’, *osm-9(ky10)* 30’ vs 120’, wild-type 0’ vs 120’, *osm- 9(ky10)* 0’ vs 120’, and wild-type 30’ vs 120’, all after three-cycle spaced-training.

**Fig 3.**
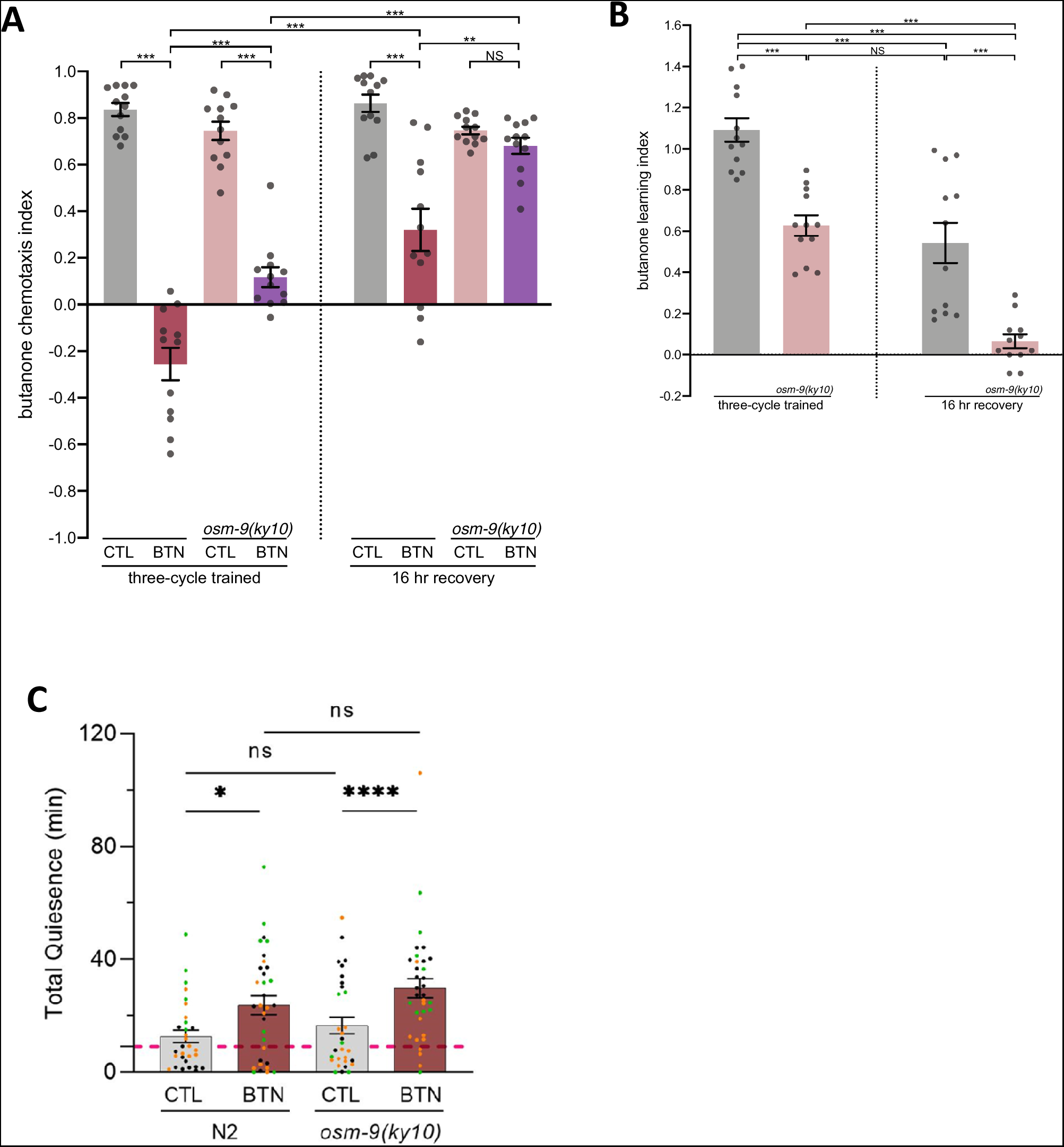
*osm-9* animals do not maintain the memory 16 hours after training. (A) Chemotaxis indices for three-cycle trained animals after 0’ vs 16 hours of recovery on food. The Kruskal-Wallis test was performed. The u-test was performed for *osm-9(ky10)* CTL vs *osm-9(ky10)* BTN, CTL 16 hr vs BTN 16 hr, *osm-9(ky10)* BTN vs *osm-9(ky10)* BTN 16hr, and BTN vs *osm-9(ky10)* BTN. The t-test was performed for CTL vs BTN, BTN 16 hr vs *osm-9(ky10)* BTN 16 hr, *osm-9(ky10)* CTL 16 hr vs *osm-9(ky10)* BTN 16 hr, and BTN vs BTN 16 hr. (B) Learning indices for three-cycle trained animals after 0’ vs 16 hours of recovery on food. The Kruskal-Wallis test was performed. The u-test was performed for wild type 16 hr vs *osm-9(ky10)* 16 hr, wild type 0’ vs wild type 16 hr, and *osm-9(ky10)* 0’ vs wild type 16 hr. The t-test was performed for wild type 0’ vs *osm-9(ky10)* 0’, *osm-9(ky10)* 0’ vs *osm-9(ky10)* 16 hr, and wild type 0’ vs *osm-9(ky10)* 16 hr. (C) Individual three cycle trained wild type or *osm-9(ky10)* animals were placed (immediately after training) into the wells of WorMotels and imaged for two hours in order to assess their quiescence as an indicator of sleep. Each datum point is one animal and the experiment was repeated three times. Quiescence is not different between wild type and *osm-9(ky10)* mutant animals after either training.

### Sleep after training is intact in *osm-9* mutant animals

Long term (16 hour long) memory consolidation was shown to require sleep in Chandra et al., 2022 [27] and so we asked if the consolidation defects in *osm-9(ky10)* mutant animals might be due to an inability to sleep. We placed 3 cycle-trained *osm-9(ky10)* and wild type animals into a WorMotel and recorded their movement for two hours after training. We found that whether the *osm-9(ky10)* animals were buffer or butanone trained, they were quiescent for the same length of time after training as the respective buffer or butanone trained wild type animals (Figure 3 C). Thus, the *osm-9(ky10)* mutants’ inability to consolidate memory is not due to their inability to sleep.

### Two independent alleles of *osm-9* show long term memory defects

We next tested additional *osm-9* alleles to understand if the *osm-9(ky10)* allele is unique or whether other alleles show the same phenotype. Having more than one allele that shows a similar phenotype can indicate that the mutant phenotype results from mutation of the gene of interest rather than a “hidden” mutation at another location in the one allele’s genome. The *osm-9* gene has fourteen exons and the *ky10* allele encodes an early stop codon at the beginning of the fourth exon. This stop codon occurs in the middle of the third ankyrin repeat (Fig 4A) and was generated in a screen for adaptation defective mutants in the Bargmann lab where it was also found to be defective for chemotaxis to AWA sensed odors such as diacetyl [37]. In a separate screen, the Sze lab isolated the *osm-9(yz6)* allele [87], which harbors an early stop codon mutation at the end of the fifth exon, 3’ of the six ankyrin repeats (Fig 4A). mRNA transcripts with early stop codons such as these tend to be degraded via the nonsense mediated decay pathway and often reveal the null phenotype. We found that the *yz6* mutant strain was unable to seek out a point source of diacetyl (Figure 4B). The *yz6* allele, like *ky10,* was defective for both one cycle and three cycle-learning, as well as the long lasting 16-hour memory that requires sleep for consolidation (Figure 4C and D).

**Fig 4.**
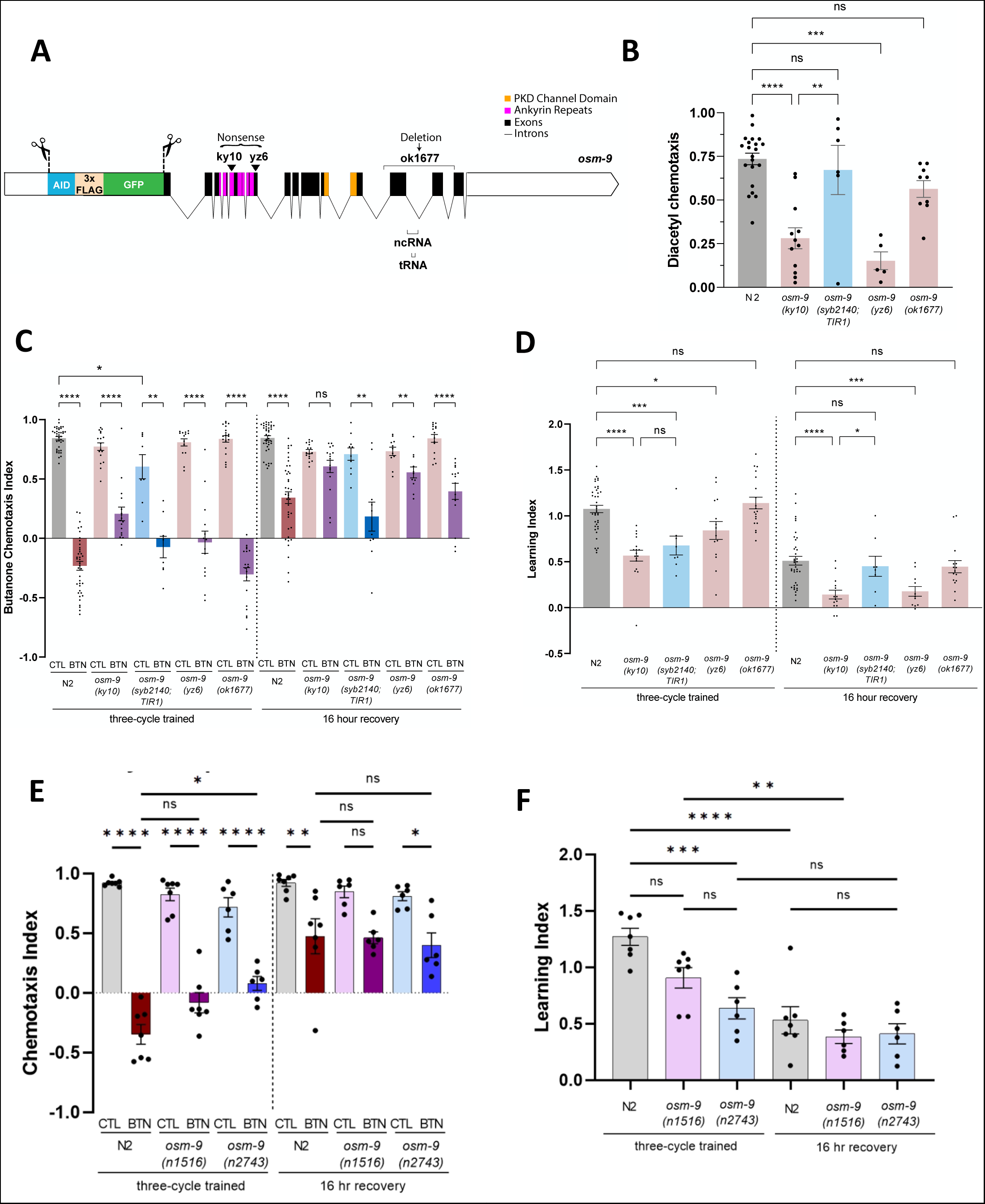
Both endogenously tagged GFP-OSM-9 *(syb214)* and *osm-9(ok1677)* show wildtype behavior while both *ky10* and *yz6* are defective for AWA-mediated chemotaxis and LTM. (A) The endogenous copy of the *osm-9* gene was edited using CRISPR to encode a fusion between OSM-9 and GFP, 3x FLAG, and the <u>a</u>uxin inducible <u>d</u>egradation signal AID at its N-terminus. This is designated *osm-9*(*syb2140*). On the right, the gene map of *osm-9* is shown. Exons are denoted by black boxes and introns are black lines connecting the exons. Red-violet indicates ankyrin repeats (six total). Orange indicates a PKD channel domain. The 5’ end white box represents an ∼1.6 kb promoter region [37]. The 3’ end white box represents an ∼3.2 kb promoter region [37]. Scale bar indicates 1000 bases. Black arrowheads indicate alleles with point mutations producing early stop codons (*ky10* and *yz6*). The bracket labeled *ok1677* denotes a deletion allele. The brackets labeled ncRNA and tRNA are predicted RNAs produced from these regions. The bracket labeled pore loop denotes the region permeable to calcium, located between the fifth and sixth transmembrane domains. (B) Chemotaxis indices for naive wild-type (N2), *osm-9(ky10), osm-9(syb2140;TIR1), osm-9(yz6),* and *osm-9(ok1677)* animals to the odor diacetyl. *osm-9*(*syb2140;TIR1*) was generated by crossing *osm-9(syb2140)* with p*eft-3*::TIR1::mRuby animals. The chemotaxis index (CI) is calculated as the number of worms at the diacetyl point source subtracted by the diluent (200-proof ethanol), divided by the total number or worms (excluding any at the origin). Wild-type (N2) is shown with grey bars, osm-9 mutants with pink bars, and *osm-9(syb2140;TIR1)* with blue bars. One-way ANOVA followed by Bonferroni correction was performed for wild-type (N2) vs *osm-9(ky10)*, *osm-9(ky10)* vs *osm-9(syb2140;TIR1)*, N2 vs *osm-9(syb2140;TIR1)*, N2 vs *osm-9(yz6)*, and N2 vs *osm- 9(ok1677)*. (C) Chemotaxis indices for three-cycle trained animals with no recovery or 16 hours recovery on food for wild-type (N2), *osm-9(ky10), osm-9(syb2140;TIR1), osm-9(yz6),* and *osm-9(ok1677)* animals to butanone. Grey and red bars are wild-type buffer- and butanone-trained animals, pink and purple are osm-9 buffer- and butanone-trained animals, and blue and navy bars are *osm-9(syb2140;TIR1*) buffer- and butanone-trained animals. Mann-Whitney u-test (refer to Figure 1 legend) was performed for three-cycle N2 CTL vs BTN, three-cycle *osm-9*(*ky10*) CTL vs *osm-9*(*ky10*) BTN, three-cycle *osm-9*(*syb2140;TIR1*) CTL vs *osm-9*(*syb2140;TIR1*) BTN, three-cycle *osm-9*(*yz6*) CTL vs *osm-9*(*yz6*) BTN, three-cycle *osm-9*(*ok1677*) CTL vs osm-9(*ok1677*) BTN, 16 hr N2 CTL vs BTN, 16 hr *osm-9*(*ky10*) CTL vs *osm-9*(*ky10*) BTN, 16 hr *osm- 9*(*syb2140;TIR1*) CTL vs *osm-9*(*syb2140;TIR1*) BTN, 16 hr *osm-9*(*yz6*) CTL vs 16 hr *osm-9*(*yz6*) BTN, 16 hr *osm-9*(*ok1677*) CTL vs *osm-9*(*ok1677*) BTN, and three-cycle N2 CTL vs three-cycle *osm-9*(*syb2140;TIR1*) CTL. (D) Learning indices for three-cycle trained with no recovery and 16 hours recovery on food for wild-type (N2), *osm-9*(*ky10*), *osm-9*(*syb2140;TIR1*), *osm-9*(*yz6*), and *osm- 9*(*ok1677*) animals. The learning index (LI) is calculated as the chemotaxis index of the BTN animals subtracted from the chemotaxis index of the CTL animals. The higher the learning index, the more the animals have learned and thus kept the BTN memory. Wild-type (N2) is shown with grey bars, osm-9 mutants with pink bars, and *osm- 9*(*syb2140;TIR1*) with blue bars. Mann-Whitney u-test (refer to Figure 1 legend) was performed for three-cycle N2 vs *osm-9*(*ky10*), three-cycle *osm-9*(*ky10*) vs *osm- 9*(*syb2140;TIR1*), three-cycle N2 vs *osm-9*(*syb2140;TIR1*), three-cycle N2 vs *osm- 9*(*yz6*), three-cycle N2 vs *osm-9*(*ok1677*), 16 hr N2 vs *osm-9*(*ky10*), 16 hr *osm-9*(*ky10*) vs *osm-9*(*syb2140;TIR1*), 16 hr N2 vs *osm-9*(*syb2140;TIR1*), 16 hr N2 vs *osm-9*(*yz6*), and 16 hr N2 vs *osm-9*(*ok1677*). (E) Chemotaxis indices for three-cycle trained animals with no recovery or 16 hours recovery on food for wild-type (N2), *osm-9(n15160*) first ankyrin repeat mutated*, or osm-9(n2743)* second ankyrin repeat mutated animals to butanone. Each datum point represents one independent assay with at least 50 animals. (F) Learning indices for three-cycle trained animals with no recovery or 16 hours recovery on food for wild-type (N2), *osm-9(n15160*) first ankyrin repeat mutated*, or osm-9(n2743)* second ankyrin repeat mutated animals to butanone.. Each datum point represents one independent assay from (E).

### Neither the C-terminus, ncRNA nor the tRNA of the *osm-9* locus are required for LTM or diacetyl chemosensation

We also examined the *ok1677* allele, which has a deletion of the twelfth, thirteenth, and part of the fourteenth exons which removes most of the C-terminal intracellular tail, and the sequences that produces a non-coding RNA and tRNA (Fig 4A). This mutant is predicted to encode an intact TRP channel because the residues missing from its C terminus are not predicted to contribute to the pore or TRP domain. The deletion likewise, leaves intact the potential 2-aminoethoxydiphenyl borate (2-APB) binding domain that was shown to serve as an agonist or antagonist in the mammalian TRPV 6 [83]. We found that the *ok1677* animals were able to chemotax to diacetyl (Figure 4B which is different from what was found in [37]) and had wild-type memory (Figure 4C and 4D), indicating that the C-terminus is not required in this learning paradigm. The tRNA and the noncoding RNA that map to the 12th intron likewise are not needed for either behavior.

### GFP tagging the N-terminus of endogenously produced OSM-9 does not perturb sleep-dependent long-term memory

We, with the help of SUNYBiotech, created a new allele of *osm-9(syb2140)* using a CRISPR-based strategy to tag the endogenous OSM-9 gene with GFP (Figure 4A). The *osm-9(syb2140);TIR1* strain was indistinguishable from the wild type in its ability to chemotax towards the AWA sensed odor, Diacetyl (Figure 4B). It showed a mild chemotaxis defect to butanone which resulted in a learning defect after three cycle (3x) training with butanone and starvation (Figure 4C and D), but this defect did not impair sleep-dependent long-term memory as this strain’s Learning Index (LI) after 16 hours of recovery (R16) was indistinguishable from the wild type (Figure 4D). We used the double, *osm-9(syb2140);peft-3::TIR1::mcherry* for most of the butanone LTM assays and since the integrated strain CA1200 (*peft-3::TIR1::mcherry)* denoted TIR1 has a mild butanone response defect, the double’s deficits in learning must be interpreted cautiously. We are in the process of repeating these studies with the single *osm- 9(syb2140)* strain.

### The point mutations of conserved glycines in either of the first two ankyrin repeats do not affect long term memory

Mutations in either of the first two of the OSM-9 six predicted ankyrin repeats (*n1562* and *n2743,* respectively) were shown to block diacetyl chemotaxis [37]. We found that animals carrying a mutation in either repeat showed nearly wild type memory acquisition and long term memory (**Figure 4** **E** and **F**). Populations carrying a mutation in the second ankyrin repeat did show a learning defect LI at t=0 as their LIs were significantly lower than wild types (**Figure 4F**). Thus, protein-protein interactions that are mediated by the first and possibly second ankyrin domains, either are dispensable for memory consolidation or do not negatively impinge upon butanone memory consolidation.

### Expressing OSM-9 from curated promoters failed to rescue the long term memory defect of the *osm-9(ky10)* mutant strain

To understand what cells *osm-9*/*TRPV5/TRPV6* is required in for wild type long term memory, we first took a molecular genetic approach. We asked whether *osm-9* transgenes under various promoters could rescue the *osm-9(ky10)* mutant strain’s defects. We were unable to rescue the *osm-9(ky10)* strain’s short-term (80 minute exposure, no food recovery) odor-learning defect [25, 51] (S1 Fig) with a partial *osm-9* cDNA under its endogenous upstream and downstream promoters (S3D Fig), which was previously shown to drive *osm-9* expression and localization in AWC and to rescue the *ky10* allele’s diacetyl chemotaxis defects [37,38 the clone is called *osm-9*::GFP5 in [37]]. We also could not achieve rescue with the cDNA driven by an AWC specific-promoter (p*ceh-36*; S3A Fig), the olfactory sensory neuron promoter (p*odr-3,* expresses in AWA, AWB, and AWC; S3B and S3C Figs), under both of its own upstream and downstream promoters [37] (S3E Fig), or a ciliated neuron promoter (*nphp-4,* [56]; S3G Fig). Nor did we observe rescue using a fosmid containing the whole *osm-9* gene (S3F Fig).

Partial rescue was achieved using the full-length *osm-9* cDNA under both the olfactory sensory neuron promoter *odr-3* and its own downstream promoter. Full rescue was only obtained when expressing the *osm-9* genomic DNA under both its upstream and downstream promoters, although the extrachromosomal array silenced in all of the transgenic lines after about 4 generations, precluding further rescue analysis (S5 Fig). The lack of rescue with curated promoters that express in AWC made us unsure of the endogenous expression pattern of OSM-9.

### Endogenous OSM-9 expression is seen in sensory neurons in the head and tail

We thus decided to examine the endogenous expression of OSM-9 using a CRISPR based strategy to tag OSM-9 with GFP. As shown in Figure 5A, we created a fusion between OSM-9 and GFP, 3x FLAG, and the auxin inducible degradation signal AID (Figure 5A). The OSM-9 expression pattern was examined in the resulting *osm- 9(syb2140)* strain (Figure 5A). We focused on the boxed regions in panel B.

**Fig 5.**
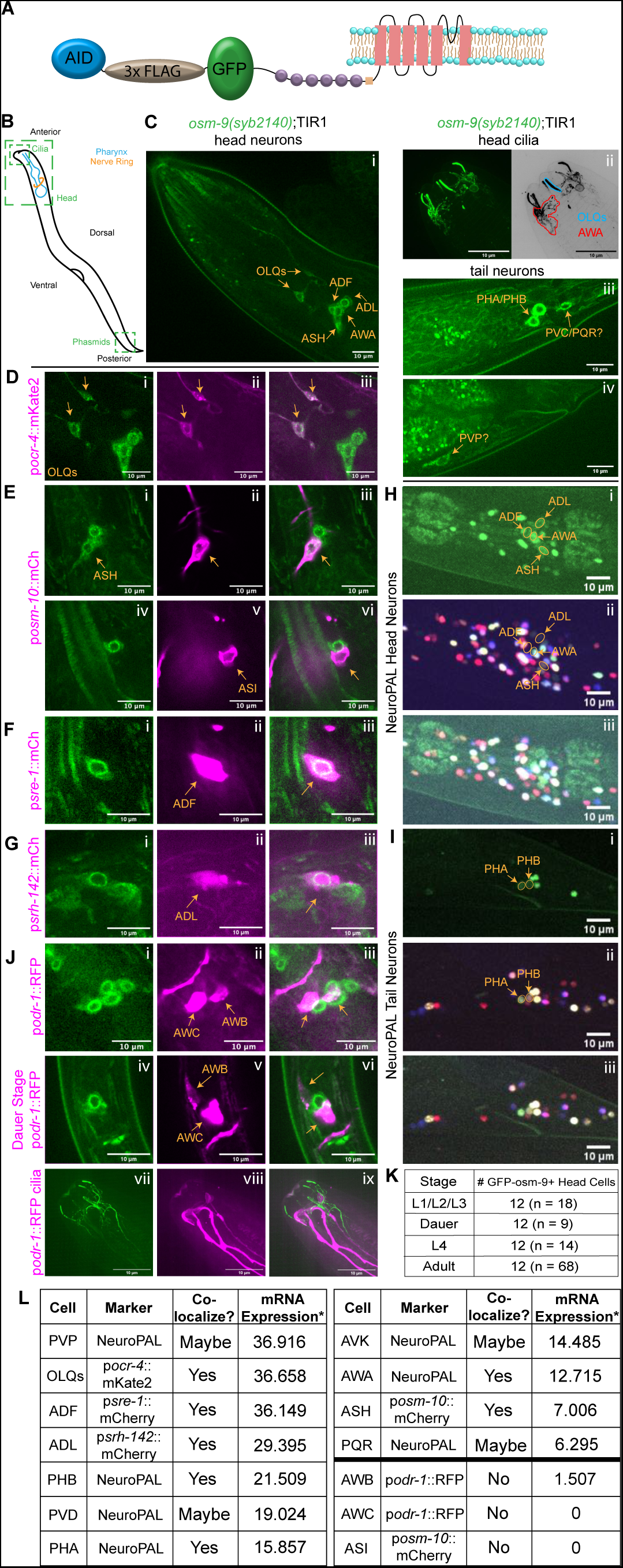
Endogenous GFP-OSM-9 is expressed in head and tail sensory neurons but not AWC. (A) The endogenous copy of the *osm-9* gene was edited using CRISPR to encode a fusion between OSM-9 and GFP, 3x FLAG, and the <u>a</u>uxin inducible <u>d</u>egradation signal AID at its N-terminus as indicated in the cartoon. (B) Schematic of C. elegans, boxes indicate regions imaged: whole head, buccal region with cilia, and tail region with phasmids. (C) Left: Visualization of endogenous GFP-tagged OSM-9 expression in the *osm- 9*(*syb2140*);p*eft-3*::TIR-1::mRuby strain in the head; cell bodies located in the anterior ganglion and lateral ganglion marked with arrows (i). Top right: In the tip of the nose, the “tree-like branching” AWA cilia (outlined in red) and the “rod-like” OLQ cilia (marked in blue) (Doroquez et al., 2014) express GFP (ii). Middle right: In the tail region’s lumbar ganglion, the phasmids PHA/PHB also express GFP (iii). A third neuron in the lumbar ganglion, possibly PVC or PQR, also expresses GFP (iii). Bottom right: Another neuron in the pre-anal ganglion, possibly PVP, also expresses GFP (iv). Anatomical regions are as defined in Yemini et al., 2021. (D) p*ocr-4*::mKate2 marks the OLQ neurons and colocalizes with GFP-OSM-9 in *osm- 9*(*syb2140*);p*eft-3*::TIR-1::mRuby (i-iii). The p*eft-3*::TIR-1::mRuby signal is very dim and not observed in the head region. (E) p*osm-10*::mCherry marks the ASH (ii and iii) and ASI (v and vi) neurons in the *osm- 9*(*syb2140*);p*eft-3*::TIR-1::mRuby strain. There is overlap between mKate2 and GFP in ASH (i-iii), but not in ASI (iv-vi). These sets of neurons were imaged from the same animal. (F) p*sre-1*::mCherry marks the neuron ADF in *osm-9*(*syb2140*) and GFP-OSM-9 colocalizes (i-iii). (G) p*srh-142*::mCherry marks the neuron ADL in *osm-9*(*syb2140*) and GFP-OSM-9 colocalizes (i-iii). (H) *osm-9*(*syb2140*);NeuroPAL animals showed colocalization between the ASH, AWA, ADL, and ADF neuronal cell bodies in the NeuroPAL channel and the GFP channel in the head (i-iii). (I) *osm-9*(*syb2140*);NeuroPAL animals showed colocalization between PHA and PHB in the NeuroPAL channel and the GFP channel in the tail (i-iii). (J) p*odr-1*::RFP marks AWC and AWB in the *osm-9*(*syb2140*);p*eft-3*::TIR-1::mRuby strain. Top: No colocalization is seen in either neuron in adult animals (i-iii). Middle: Animals in day 1 dauer stage do not show colocalization between AWC, AWB, and GFP (iv-vi). Bottom: In the tip of the nose, AWC and AWB “wing-like” cilia do not colocalize with GFP cilia (vii-ix). (K) The larval stages, dauers, L4s, and adult animals all show 12 GFP-expressing neurons total in the head. (L) Summary of endogenous OSM-9 *osm-9*(*syb2140*) expression. *CenGen data (https://cengen.shinyapps.io/SCeNGEA/).

In the head, we found that OSM-9::GFP is expressed in six sets of neurons (12 in total) whose cell bodies are close to the anterior bulb of the pharynx (Figure 5 Ci). The endings of the head neurons appeared in the buccal area around the lips of the worm. Using super-resolution (SORA) microscopy and comparing the structures to the EM reconstructions published in Doroquez et al., 2014 [57], we were able to identify the ciliated rod-like structures of the OLQ neurons and the “tree-like branching” AWA cilia (Figure 5 Cii). The finger-like ADL sensory endings were faintly visible (data not shown). In the tail region, we noted three pairs of neuronal cell bodies in the lumbar ganglion (Figure 5 Ciii as described in Yemini et al., 2021 [58]) and two pairs in the pre- anal ganglion. The four neurons in the anterior ganglion were verified as the OLQs because their cell bodies co-localized with mKate2 expression driven by the *ocr-4* promoter (Figure 5 D i-iii).

GFP-OSM-9 expression was identified in the ASH cell body because it colocalized with mCherry driven by the *osm-10* promoter, which is expressed in the dorsally located ASH (Figure 5 E i-iii). This promoter also allowed us to rule out OSM-9 expression in ASI as the ASI cell body is located ventral to that of ASH and this ventral *posm-10*::mCherry positive cell body did not colocalize with GFP-OSM-9 (Figure 5E iv-vi).

The ADF cell body was marked by its expression of the mCherry reporter under the control of the *sre-1* promoter and GFP-OSM-9 colocalized with it (Figure 5 F i-iii). Thus, OSM-9 is expressed in ADF. In a similar manner, we used mCherry expression from the *srh-142* promoter to identify OSM-9 expression in ADL (Figure 5 G i-iii). By creating a double with *osm-9(syb2140)* and the recently developed NueroPal strain (*otls670*) [58], we were able to verify that OSM-9 is expressed in ASH, ADF and ADL as well as AWA (Figure 5 H i-iii). Thus, we corroborated its expression in AWA using both the cilia morphology (Figure 5 Cii) and colocalization of expression with known promoter expression (Figure 5 H i-iii).

In the tail region, we found that OSM-9::GFP is expressed in the cell bodies of three pairs of neurons near the lumbar ganglion in the tail which we identified as PHAR/L and PHBR/L and possibly PQRR/L. In the pre-anal ganglion, we noticed a cell body that is most likely PVP (Figure 5 I i-iii).

### Endogenous GFP-OSM-9 expression is not observed in the butanone responsive AWC neuron

Unexpectedly, though we and others had posited that OSM-9 is expressed in and may function in AWC to permit odor learning, GFP-OSM-9 was not observed in the cell body or the sensory endings of the AWC sensory neuron (Figure 5 J i-ix). We used expression of RFP from the *odr-1* promoter to mark the cell bodies and cilia of the AWC and AWB neurons (noted as the *gcy-10* promoter in Yu et al. 1997 [59]) and found that GFP-OSM-9 was not colocalized with RFP at any stage of development including dauer animals (Figure 5 J i-ix). Likewise, though the wing-like sensory endings of AWC and AWB were clearly marked with RFP, we saw no evidence of co-localized GFP-OSM-9 (Figure 5 J iiv-ix). Consistent with our inability to observe GFP-OSM-9 in AWC or AWB, the CenGen data set indicates that they failed to see evidence of *osm-9* mRNA expression in AWC and its expression in AWB was extremely low (Figure 5 K) [68]. Though the expression of GFP-OSM-9 may, indeed, be extremely low in the AWB and AWC neurons and we may need to use immunological techniques to amplify the signal, we favor the new hypothesis that OSM-9 functions in the circuit to allow for olfactory learning and consolidation.

As might be expected for a membrane protein, GFP-OSM-9 staining is excluded from the nucleus of the amphid neurons and travels in packets anteriorly from the cell body to cilia (R.C., personal observations). The ambiguity in its expression in PVP, PVD, AVK and PQR could arise from concentration of the GFP-OSM-9 signal within structures such as sensory endings and neurites that are distal from the cell bodies which we used for NeuroPAL verification.

### Butanone exposure does not alter GFP-OSM-9 expression in the wild type genetic background

Our previous work had placed OSM-9 downstream factors that act in the AWC nucleus to promote learning. We had shown that loss of OSM-9 permits chemotaxis in animals that expressed gain-of-function forms of the nuclear factors EGL-4 and HPL-2 [49, 50, 51]. We interpreted this to mean that the nuclear factors, acting either in AWC or in other neurons, might promote OSM-9 expression and over-expressed OSM-9 would decrease odor responsiveness.

If this hypothesis were correct, we might expect that butanone training would increase the numbers of cells that express GFP-OSM-9 and/or increase expression in the cells we identified in naive animals. We looked exhaustively throughout the head of butanone-trained animals but were unable to identify any new cell bodies or changes in overall expression patterns after training.

Thus, we decided to quantify GFP-OSM-9 expression level to see if we could detect more subtle changes. We focused on GFP-OSM-9 expression in the animal’s nose tip, which harbors the AWA, OLQ sensory endings and cilia and the head region where we observed the AWA, ADF, ADL and ASH cell bodies (Figure 6A). We quantified the levels of GFP expression in the cell bodies and cilia of animals that had either been exposed to buffer or butanone for: 80’ (labeled 1X), three cycles of 80’ interleaved with 30’ of food (3X) or three cycles, 3X followed by 16 hours of recovery on plates with food (16 hr. Rec.) (Figure 6B). We found that the levels of GFP-OSM-9 in either the cell body or the cilia were not changed by treatment (Figure 6B). The levels of GFP-OSM-9 did increase in the cilia after 16 hours of recovery but these increases were not specific to odor treatment and thus may be due to age-dependent increases in cilia size.

**Fig 6.**
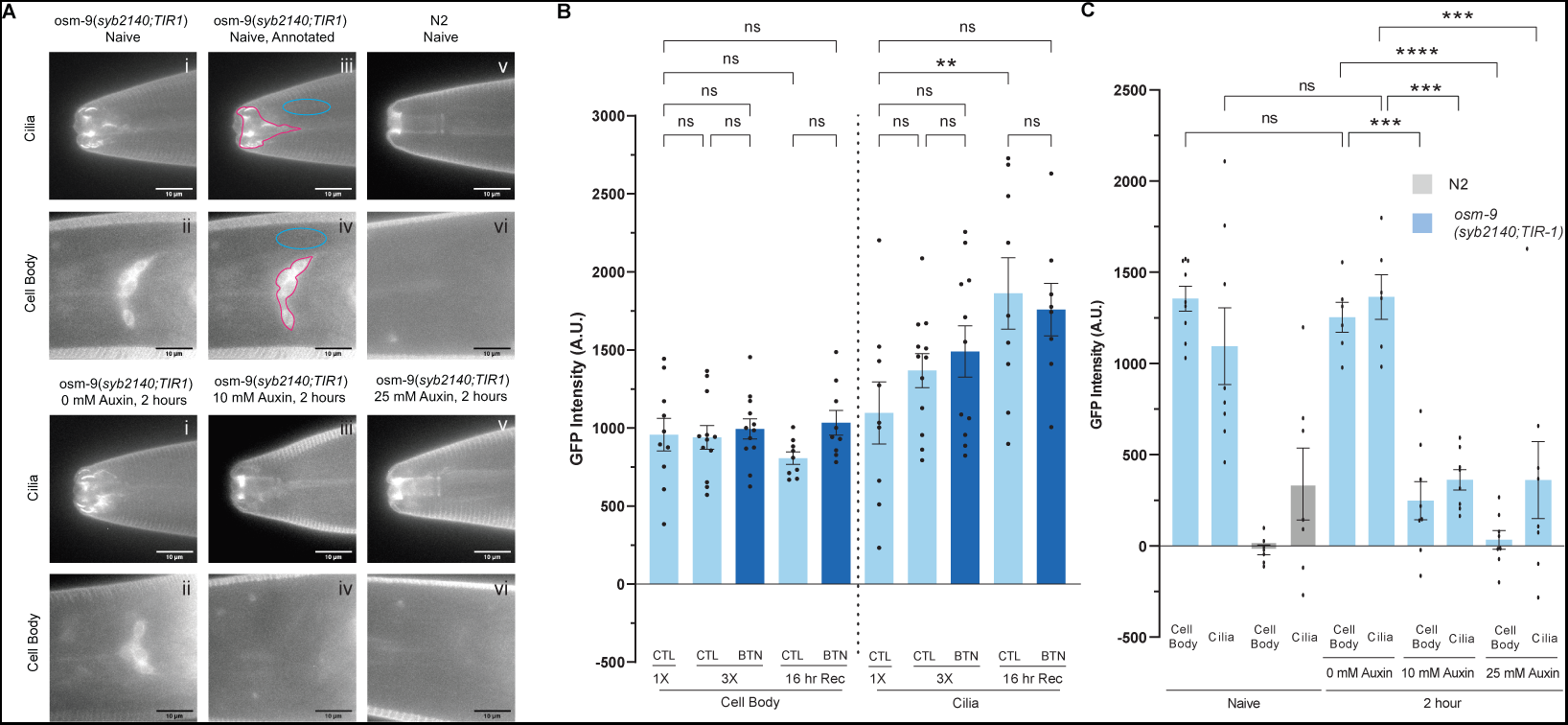
Two Hours of Auxin-Induced Degradation Reduces Endogenous GFP-OSM-9. (A) Representative images of *osm-9*(*syb2140;TIR1*) GFP expression in the cell body and cilia before auxin treatment (Naive) (i-ii), after 2 hours of 0 mM auxin treatment (vii-viii), after 2 hours of 10 mM auxin treatment (ix-x), and after 2 hours of 25 mM auxin treatment (xi-xii). TIR1 is the substrate recognition component of the auxin inducible E3 ligase and it is expressed from the ubiquitous promoter *eft-3*. Once TIR1 binds auxin, it induces degradation of target proteins that contain the AID. Example region analyzed denoted by magenta outline and region used for background subtraction denoted by blue outline (iii-iv). (B) GFP intensities of cell bodies and cilia of *osm-9*(*syb2140;TIR1*) animals after one cycle (1X) training, three-cycle (3X) training, and 16 hour recovery on food after 3X training shows a significant increase in GFP intensity after 16 hours recovery in control animals. One-way ANOVA followed by Bonferroni correction was performed for 1X CTL cell body vs 3X CTL cell body, 3X CTL cell body vs 3X BTN cell body, 1X CTL cell body vs 3X BTN cell body, 1X CTL cell body vs 16 hr Recovery CTL cell body, 1X CTL cell body vs 16 hr Recovery BTN cell body, 16 hr Recovery CTL cell body vs 16 hr Recovery BTN cell body, 1X CTL cilia vs 3X CTL cilia, 3X CTL cilia vs 3X BTN cilia, 1X CTL cilia vs 3X BTN cilia, 1X CTL cilia vs 16 hr Recovery CTL cilia, 1X CTL cilia vs 16 hr Recovery BTN cilia, and 16 hr Recovery CTL cilia vs 16 hr Recovery BTN cilia. (C) GFP intensities of cell bodies and cilia in naive *osm-9*(*syb2140;TIR1*) and N2, as well as *osm-9*(*syb2140;TIR1*) animals incubated in 0, 10, or 25 mM K-NAA synthetic auxin shows significant loss of OSM-9 after 2 hour auxin incubation. One-way ANOVA followed by Bonferroni correction was performed for Naive cell body vs 0 mM Auxin cell body, 0 mM Auxin cell body vs 10 mM Auxin cell body, 0 mM Auxin cell body vs 25 mM Auxin cell body, Naive cilia vs 0 mM Auxin cilia, 0 mM Auxin cilia vs 10 mM Auxin cilia, and 0 mM Auxin cilia vs 25 mM Auxin cilia.

### Auxin degradation system reduces endogenous GFP-OSM-9

In creating the GFP-tagged allele of *osm-9*, we added sequences that encode for an auxin-dependent E3-ligase recognition site [60]. In this way, we should be able to reduce the levels of OSM-9 by expressing the E3 Ligase TIR1 and treating the animals with auxin. We expressed TIR1 from the ubiquitous *eft-3* promoter and introduced this integrated transgene into *osm-9(syb2140)* by crossing the strains. Naive *osm- 9(syb2140);*p*eft-3::*TIR1 animals that are straight off their growth plates show cell body and cilia expression that is not different from that of *osm-9(syb2140);*N2 animals (Figure 6C). The level of GFP-OSM-9 is decreased to less than one half of the naïve after two hours of incubation with 10mM auxin. Incubation with 25mM auxin further reduced expression such that it was undetectable in the cell bodies but it never completely disappeared from the cilia. The cilia were notably different from the N2 staining pattern, which shows a bright buccal cavity autofluorescence. Thus, this tool may need to be supplemented with other ways to remove OSM-9 in order to better assess its role in sleep-dependent olfactory memory.

### Chronic, ZIF-1::GFP-nanobody mediated degradation of GFP-OSM-9 in ciliated neurons blocks long-term memory consolidation

In order to chronically deplete OSM-9 and thus perhaps phenocopy the null, we decided to utilize the ZIF-1 toolkit for degradation of GFP-tagged proteins [88]. This takes advantage of the *C. elegans* SOC-box adapter protein ZIF-1 that is fused to a nanobody that recognizes GFP to bring the cullin2 E3 ubiquitin ligase to a GFP-tagged protein ([88] and Figure 7A). Wang et al., 2017 developed the p*osm-6*::ZIF-1 (OD2772) strain that expresses the ZIF-1-nanobody fusion from a promoter that is expressed in all ciliated neurons along with an mCherryHistone H2B to mark the nuclei of cells that express ZIF-1-nanobody. We crossed *osm-9(syb2140)* into this strain and examined the homozygous doubles for GFP-OSM-9 expression. In all animals, no matter tehir treatment, we observed OLQ and AWA sensory endings. In very preliminary studies, we found that approximately one half of the animals examined (only 4) showed no expression in ADL, ADF and ASH and new expression in the non-ciliated AVK neuron.

**Fig 7.**
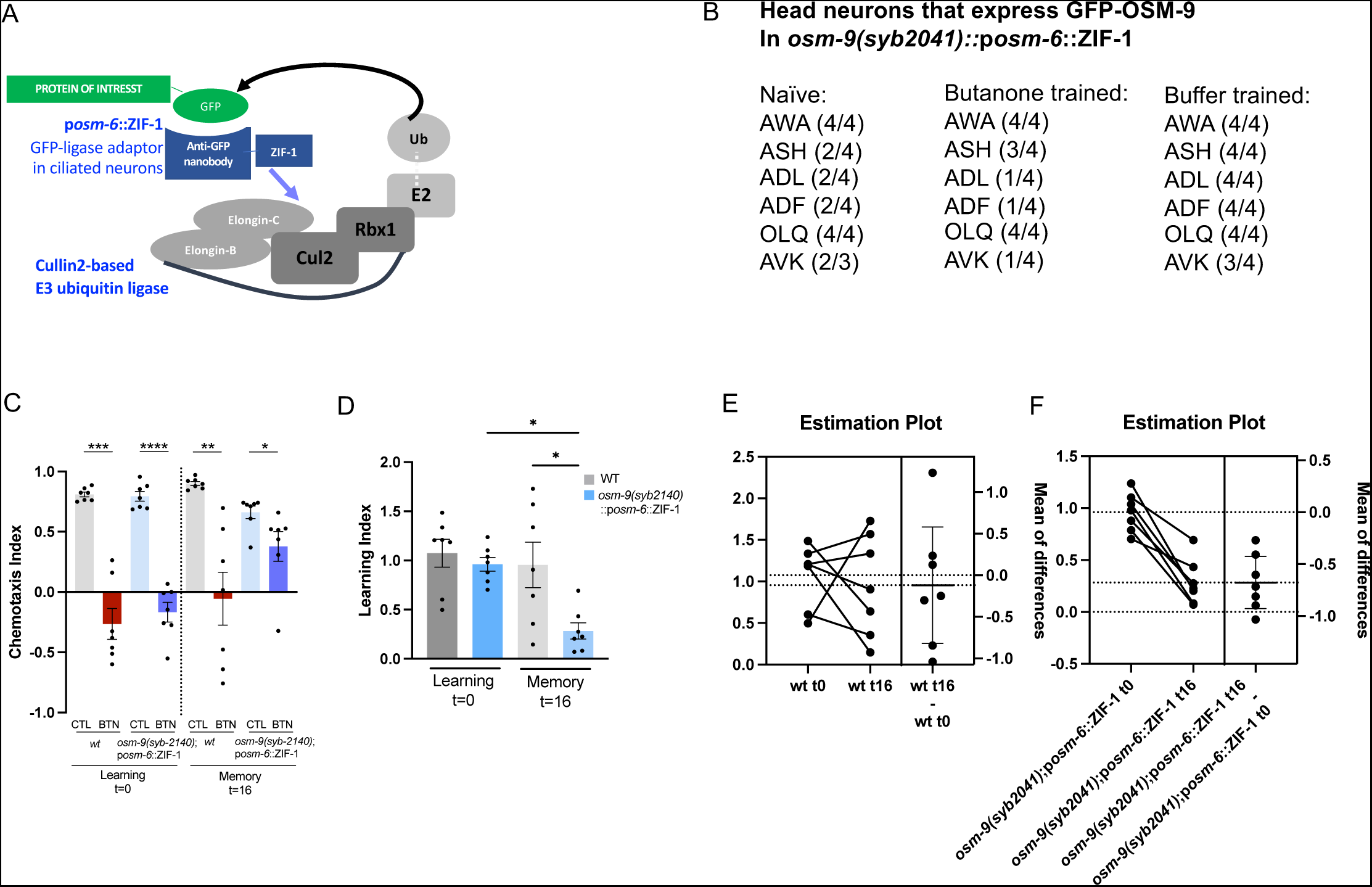
ZIF-1-nanobody Mediated Degradation System Expressed Throughout Development from the Two-Fold Embryo Stage Reduces Endogenous GFP-OSM-9 in ADL, ADF but not AWA in Naïve Animals and Reveals Long-Term Memory Defects. (A) ZIF-1 mediated degradation of target GFP-tagged proteins [88].. The *C. elegans* SOCs box ZIF-1 protein was appended to an anti-GFP nanobody which brings the ubiquitination ligase complex in to contact with the protein target which is ubiquitinated and degraded by the cellular proteasome. Schematic is adapted from Wang et al., 2017. Endogenously expressed GFP tagged OSM-9 is targeted by expression of ZIF-1 from the *osm-6* promoter which is expressed in all ciliated neurons beginning at the 2- fold stage in the embryo in many ciliated neurons [89: Supplemental information S3]. (B) Preliminary analysis of cells in the *osm-9(syb2140); posm-6*::ZIF- 1::SL2::mCherryH2B (*syb2140;OD2772*) that express GFP in four day one adults. Animals were imaged at 400X on the WS2 spinning disk confocal. (C) Chemotaxis indices for three-cycle trained animals with no recovery or 16 hours recovery on food for wild-type (N2), *osm-9(syb2140); posm-6*::ZIF-1::SL2::mCherryH2B expressing animals to butanone. Each datum point represents one independent assay with at least 50 animals. (D) Learning indices for three-cycle trained animals with no recovery or 16 hours recovery on food for wild-type (N2), *osm-9(syb2140); posm-6*::ZIF-1::SL2::mCherryH2B carrying animals to butanone.. Each datum point represents one independent assay from (C).

Expression in AWA was unchanged (Figure 7B). When we examined four butanone trained animals, we found that one quarter showed ADF and ADL expression, three quarters ASAH and all showed AWA expression and one showed expression in AVK. Buffer trained animals all showed the same expression pattern as *osm-9(syb2140)* in the wild type background except it was a bit fainter. Only animals that showed mCherry stained nuclei were examined and so if the integrated transgene had been silenced, it would have had to have happened in less than the half-life of mCherry (data not shown).

When we examined the ability of the *osm-9(syb2140);*p*osm-6::*ZIF-1 strain to acquire and consolidate memory, we found that it was able to learn as well as wildtypes (Figure 7 C, D t=0) but it failed to keep the memory (Figure 7 C, D t=16). Each datum point is one day with >100 animals in the assay. The amount of memory that is lost in each strain can better be appreciated by looking at the change in LI over 16 hours (Figure 7 E,F). Thus, a very preliminary understanding is that since memory consolidation is lost in animals that express wildtype levels of OSM-9, OSM-9 expression in AWA is either not sufficient to block the negative effects of lack of expression in ADL, ADF and ASH or the negative effect of expression in AVK or loss of OSM-9 in ADL, ADF and possibly ASH leads to lack of consolidation because the activity of these cells is compromised and their activity is required for memory consolidation.

## Discussion

Transient receptor potential (TRP) channels are best understood in their role as sensors of harmful environmental stimuli, yet their role in regulating neural plasticity - a fundamental process in the brain that governs information storage and adaptation to environmental cues is more enigmatic. We found that the *osm-9* TRPV channel, which has a known role in short-term olfactory learning [25] is also required for learning after spaced training and possibly for sleep-dependent consolidation of that memory such that it is retained for at least 16 hours. Thus, wild type activity of the *osm-9* TRPV channel is key for long-term memory but the mechanism by which it acts remains a mystery.

### OSM-9 may act downstream of nuclear factors to promote three cycle learning

*osm-9* mutant animals had previously been shown to fail to acquire odor memory after one cycle of training, and here we show that they are also impaired for memory after three-cycle spaced training. The mechanism by which animals learn after one and three cycles of training may be distinct. In the nucleus, genes including the *egl-4* PKG and *hpl-2* HP1a homolog modify chromatin and could either activate genes upstream of *osm-9* to promote one-cycle induced short-term memory [49, 51] or repress the repressors of OSM-9 function. However, these factors have not been assessed for their roles in three-cycle induced long-term memory. Thus, it remains to be seen if *osm-9* acts downstream of these factors in three-cycle induced, long-term memory. If it does act downstream, they or the genes they regulate may not directly regulate OSM-9 protein levels as we have not observed any differences in *osm-9* gene expression after either one or three cycle training in the wild type background. Instead, their gene targets may be direct or indirect activators of OSM-9 function and since these nuclear factors act in the AWC nucleus [49, 51], they may affect the cells that do express OSM-9 perhaps by producing soluble ligands that control channel open probability.

Loss of *osm-9* could also suppress the constitutively “trained” phenotype of the gain-of-function EGL-4 or HPL-2 if losing OSM-9 allows the neurons that typically express OSM-9 to now respond to butanone. The other newly butanone responsive neurons would stand in for the silenced AWC neurons and allow animals to respond to butanone. These neurons may, however, not be configured properly to allow for memory formation or consolidation.

### OSM-9 may act in sensory neurons that detect the unconditioned stimulus - lack of food

The channel is most likely to act in the sensory endings or the cell body of sensory neurons that normally are not required for butanone odor-taxis since ablation of AWA, ADL, ADF or ASH does not affect butanone odor-taxis ([61] and Chantal Brueggeman, data not shown). OSM-9 may be required for the animal to sense the lack of food and in that way provide cues for the unconditioned stimulus. The *osm-9(ky10)* mutants have foraging defects in that they remain on the portions of the plate that has no food (personal observations: KB, JMM, MFS and NDL), perhaps because they do not sense signals that normally accompany the absence of food. One such signal is the ascaroside cues emitted by nematodes that allow them to sense population density. These pheromones are sensed by ADL and ADF responds to *E. coli* [62] and thus could sense its absence. The OLQ neurons are mechanosensory neurons with sensory endings in the inside of the buccal cavity that are required for response to light nose touch and the lawn of E. coli [63] and animals that do not have these neurons may not be able to sense the absence of food. They may thus be impaired for their ability to integrate the unconditioned and conditioned stimuli.

The understanding that OSM-9 acts non-cell autonomously may allow us to interpret our finding that chronic loss of OSM-9 from neurons other than AWA and the OLQs in our *posm-6*::ZIF-1 mediated degradation system leads to loss of memory consolidation. If loss of OSM-9 from ADL, ADF and or ASH makes these cells either unable to contribute to the unconditioned stimulus (starvation) or more active in responding to butanone, the result would be continued attraction to the odor even after multiple cycles of training.

### OSM-9 may act in nociceptive neurons to promote a negative internal state in response to butanone

Our work has revealed that *osm-9* is expressed in the ADF and ASH sensory neurons which mediate repulsion [68] and ASH can be stimulated by the levels of butanone that we use in our training paradigm [58]. Thus, It is possible that *osm-9* is required in these neurons to engage an aversive brain state that drives memory consolidation of the odor. Conversely, it could be required in repulsive sensory neurons for the animal to understand that it is a negative cue, and subsequently its interneuron expression is necessary for integration and consolidation and memory of these negative cues.

### OSM-9 may detect the coincidence of two stimuli and thus drive learning by integrating the sensation of butanone and an unpleasant state

Animals must respond to and learn from their ever-changing surroundings to ensure survival. TRP channels emerged early in eukaryotic evolution to allow sensation of the environment. Over the course of evolution, they became coincidence detectors that sense a range of mechanical to chemical cues, which are vital for learning and memory [69]. Originally thought of as only having peripheral nerve expression, TRPV channel proteins are also found in the central nervous system throughout the brain.

TRPV channels in the sensory neurons may respond to cues such as heat and protons, and may subsequently also drive learning and long-term memory of those cues in the brain [69]. The evolution of TRP channels could have potentially driven survival through ambient sensation and simultaneous or subsequent triggering of memory formation.

### The evolution of TRP channels from purely sensory channels to coincidence detectors required for learning and memory?

TRP channels are found throughout the domain Eukaryota, in the animal, fungi, and protist kingdoms, meaning that they evolved before multicellularity. TRPY1 is found in the single-celled fungus, *S. cerevisiae* and TRPA, C, M, L, and V channels, are found in the single-celled protist choanoflagellates [69], as well as in multicellular, bilaterally-symmetric animals, including protostome ecdysozoan invertebrates such as *Drosophila* and *C. elegans*, and deuterostome chordate vertebrates, including zebrafish, whale and Australian ghost sharks, rodents, and humans (http://ncbi.blast.gov). TRPNs are found in *Hydra magnipapillata,* which is the last common ancestor of Cnidaria and Bilateria.

Fourteen TRPA and TRPML channels were found in the sponge *A. queenslandica*, showing a diverse conservation of TRP channels since sponges are the oldest living metazoan phyletic lineage, diverging from other metazoans ∼600 million years ago [69]. TRPA1 was found to have evolved from a chemical sensor in a common bilaterian ancestor to invertebrates and vertebrates, conserved across ∼500 million years of evolution [70]. TRP channels themselves are likely even older than the nociceptive family of sensors, since fungi branched off from other life during the Proterozoic Era, about 1.5 billion years ago [71]. These early TRP channels evolved the ability to sense multiple stimuli and drive distinct responses to ensure organismal survival. For example, the TRPA1 channel likely evolved through adaptive evolution via amino acid changes in response to the environment, explaining the temperature and compound sensitivities observed in different mammalian species [69,72–74]. The TRP channel predecessors were not neuronally expressed, but as animals became more evolved, such as through development of a neural net or cephalization, TRP channels evolved neuronal roles such as sensory perception and polymodality. However, the distinct functions of TRP channels in higher level processes such as plasticity, learning, and memory are still being uncovered.

### Mechanisms of activation of TRPV 5/ 6 channels

Since TRPV channels are required for acute sensation and in plasticity, considered short and long-term processes, respectively, then how does its function and regulation differ between them? Distinct from the TRPV1-4 channels, TRPV5 and TRPV6 are highly-selective for calcium and can mediate calcium signaling through constitutive opening at physiologic membrane potentials, possibly without the need for ligand activation. Potentially, the TRPV5 and 6 ortholog OSM-9 may also be constitutively open and may mediate the calcium homeostasis necessary to drive long-term memory formation [75]. The calcium-binding, calmodulin-like domain is preserved in the *osm-9(ok1677)* allele that shows wild-type functions. In the future, we would like to test the hypothesis that OSM-9 requires this domain to promote learning and memory. Since Ca^2+^ signaling is important for synaptic plasticity, it could be that TRPV proteins are specifically required for regulating intracellular calcium to promote plasticity and thus learning and memory.

### TRP channels as drug targets in memory disorders

Besides being a current focus for chronic pain treatment (e.g. through topical capsaicin), the TRPVs are also emerging as a new drug target for diseases affecting neural plasticity. Indeed, a vast array of neurological diseases, including Alzheimer’s disease [76], Parkinson’s disease, schizophrenia [77–79], epilepsy, and Charcot-Marie-Tooth disease [80], as well as sense disorders (e.g. anosmia) [81] result from or are linked to defects in TRP function, with affected individuals exhibiting decreased neural plasticity. It has also been shown that TRPV1 could be a target for cognitive decline in Alzheimer’s patients. The levels of the TRPV1 agonist anandamide were found to negatively correlate with amyloid-β (Aβ) peptide levels in AD brains [82]. Additionally, application of the TRPV1 agonist capsaicin restores proper gamma oscillations to hippocampal slices that have disrupted gamma oscillations due to over expressed Aβ.

The efficacy of capsaicin requires TRPV1 expression as it has no effect on slices from TRPV1 knockout mice [83]. Rat models of biliary cirrhosis, which induces memory impairment, have decreased memory impairment when treated with capsaicin, which also increased TRPV1 and CREB mRNA in the CA1 area of the hippocampus [84].

Recent research even suggests the use of TRPV1 agonists as therapeutics to aid in ameliorating the effects of schizophrenia since TRPV1 is linked to pain and cognitive defects in schizophrenic patients [79]. This could indicate that TRPV1-mediated plasticity defects may underlie some of these neurological disorders.

Performing *in vivo* studies to understand how TRPV proteins regulate proper neurological function may uncover a role in learning and memory and could potentially help us to understand what happens in the TRPV1-mediated disease states. We aimed to elucidate how TRP protein functions at the molecular level to promote plasticity and memory formation. Understanding the molecular mechanisms and dynamics by which animals learn to ignore non-beneficial stimuli through TRP-mediated neural plasticity may be of importance to understanding not only how animals learn and form memories, but may also give insights into the etiology of TRP-mediated neurological diseases.

## Materials and methods

### Strains and genetics

Strains were maintained by standard methods [85] and were grown on either 5.5 cm or 10 cm Petri dishes with NGM media seeded with OP50 *E. coli*. All strains were grown at and assayed at 20°C. The wild-type strain is the *C. elegans* N2 Bristol variant. Mutants used in the study include *osm-9(ky10) IV* [25]*, osm-9(ok1677) IV, osm-9(yz6) IV*, and the SUNY BioTech-created *osm-9(syb2041).* All rescue experiments (Fig 1, S2 Fig, S4 Fig) were performed in the *osm-9(ky10) IV* background. The transgenic lines used for the rescue experiments are JZ1967 and JZ1968 [*KBC41/podr-3::osm-9::UTR* (50 ng/μl), *punc-122::mCherry* (20 ng/μl)], (Fig 1); JZ2034, JZ2035, and JZ2036 [*KBC56/posm-9::osm-9::UTR* (50 ng/μl), *pCFJ90/pmyo-2::mCherry* (2.5 ng/μl), *pSMdelta* (100 ng/μl)], (S2 Fig); JZ1655 and JZ1656 [*KBC8/pceh-36::GFP::osm-9* (100 ng/μl), *punc-122::GFP* (20 ng/μl)], (S4A Fig); JZ1621 and JZ1622 [*pJK21/podr-3::GFP::osm-9* (50 ng/μl), *punc-122::mCherry* (20ng/μl), ScaI-digested N2 DNA (30 ng/μl)], (S4B Fig); JZ1819 and JZ1874 [*KBC24/podr-3::osm-9* (50 ng/μl), *pCFJ104/pmyo-3::mCherry* (5 ng/μl)], (S4C Fig); JZ1940 and JZ1941 [fosmid WRM065bE12 (from Source BioScience, 5 ng/μl), *punc-122::GFP* (20 ng/μl), *pstr-2::GCaMP3::mCherry* (25 ng/μl)], (S4D Fig); lines containing *osm-9::GFP5* (25 ng/μl), *punc-122::mCherry* (20 ng/μl), (S4E Fig); lines containing *KBC42/posm-9::osm-9::UTR* (50 ng/μl), *punc-122::mCherry* (20 ng/μl) or *KBC42/posm-9::osm-9::UTR* (10 ng/μl), *pCFJ90/pmyo-2::mcherry* (2.5 ng/μl), *pSMdelta* (140 ng/μl), (S4F Fig); JZ1905 and JZ1906 [*KBC38/pnphp-4::osm-9::GFP* (5 ng/μl), *punc-122::mCherry* (20 ng/μl); *osm-9(ky10) IV* gDNA (8 ng/μl); *ceh-36::mCherry* (5 ng/μl)], (S4G Fig). ScaI-digested N2 DNA, *osm-9(ky10) IV* gDNA, or *pSMdelta* (empty vector) were used to bring up the total injection concentration to ensure transgenesis [86]. JZ designations are L’Etoile lab strain names. *osm-9(syb2140)* will be deposited in the CGC.

### Molecular Biology

Constructs were made using standard molecular techniques. All constructs made for this paper are available from Addgene. The construct *pJK21/ podr-3::osm- 9SS::GFP::osm-9cDNA* was made by replacing the *pstr-2* promoter in *pJK17/pstr-2::osm- 9SS::GFP::osm-9cDNA*, digested with NotI and AscI restriction enzymes, with the subcloned *podr-3* fragment from *pJK15/podr-3::osm-9SS::GFP::osm-9cDNA*, using NotI and AscI. *KBC8/pceh-36::osm-9SS::GFP::osm-9cDNA* was made by amplifying *pceh-36* with primers KLB30 (TAACGGCCGGCCTCATCAGCCTACACGTCATC) and KLB31 (ACATAGGCGCGCCCGCAAATGGGCGGAGGGT) to add FseI and AscI sites on and digesting *pAW1/podr-3::osm-9SS::GFP::osm-9cDNA* with FseI and AscI to remove *podr- 3* and subclone the *pceh-36* fragment in. *KBC41/podr-3::osm-9cDNA::UTR* was made by digesting *KBC24/podr-3::osm-9cDNA* with NheI and NcoI and subcloning in the *osm-9* downstream promoter (∼3.2kb), amplified off of phenol-extracted followed by ethanol precipitated whole-worm gDNA (Phenol pH 8 from Sigma Aldrich), using the KLB161 (AGACGCTAGCGAACTTTTTTCTTCTAATTTTTTGA) and KLB162 (AACTCCATGGTTAGGTACATTTAAGGTCGATC) primers, adding on the NheI and NcoI sites. *KBC24/podr-3::osm-9cDNA* was made by digesting *KBC23/podr-3* with KpnI and NheI and subcloning in *osm-9* cDNA, amplified from whole-worm cDNA (a kind gift from Maria Gallegos) using primers KLB25 (AACTGGTACCTCATTCGCTTTTGTCATTTGTC) and KLB90 (AGACGCTAGCATGGGCGGTGGAAGTTCG), which add on the KpnI and NheI sites. *KBC42/ posm-9::osm-9cDNA::UTR* was made by digesting *KBC41/podr-3::osm- 9cDNA::UTR* with NotI and XmaI and subcloning in the upstream *osm-9* promoter (∼1.6 kb), amplified off of phenol-extracted followed by ethanol precipitated whole-worm gDNA (Phenol pH 8 from Sigma Aldrich), using the primers KLB163 (AGACGCGGCCGCCGCGGGAGTACTTTACGGG) and KLB164 (AACTCCCGGGGTTTGGTTTCTGAAAAAATTGG), adding on NotI and XmaI sites. *KBC38*/*pnphp-4::GFP::osm-9* was made by digesting *KBC26/podr-3::osm-9cDNA::GFP* with NotI and NruI and subcloning in the *nphp-4* promoter amplified from *pAK59/pnphp-4::GFP::npp-19* (a kind gift from Piali Sengupta) with primer KLB127 (AGACGCGGCCGCCAACATTATTAATCACTGCAAC) and KLB128 (AACTTCGCGAACTTCCACCGCCCATCTCATTTTTCGAGACTTTGTTA), adding on NotI and NruI sites. KBC56 was made by the following instructions: osm-9 gDNA was amplified with the primers KLB289 (GTTGTTTACCTTTTATGTTCATCCG) and KLB290 (AAATTTTCTACTGCCTGGTATCAAA) off of phenol-extracted followed by ethanol precipitated whole-worm gDNA (Phenol pH 8 from Sigma Aldrich). The *osm-9* gDNA fragment plus extra upstream and downstream homologous sequence was amplified off of the gDNA amplified with KLB289 and KLB290, using the primers KLB298 (GTTGTTTACCTTTTATGTTCAT) and KLB299 (AAAATGATCCACATAAAATTTTCTACTGCCTGGTAT). The *osm-9* upstream partial fragment (∼1.2 kb) was amplified using the primers KLB296 (TGATTACGCCAAGCTTCGCGGGAGTACTTTACGG) and KLB297 (TAAAAGGTAAACAACTAGTTTTTAGTACATGAAATAATT) off of the template KBC42/*posm-9::osm-9cDNA::UTR*, and the downstream promoter partial fragment was amplified with KLB300 (TATGTGGATCATTTTTGTCTC) and KLB301 (CCGCGCATGCAAGCTTTTAGGTACATTTAAGGTCGAT) off of KBC42/ *posm-9::osm-9cDNA::UTR*. The three fragments were then recombined in the *pSM* plasmid (linearized with HindIII) using the Takara Infusion HD Cloning kit. The *pSM* plasmid was from Steve McCarroll via Cori Bargmann.

### Short-term memory chemotaxis assay

We utilized the chemotaxis assay from Bargmann et al., 1993 [45]. About 4-5 Larval stage 4 (L4) animals were put onto NGM plates seeded with OP50 *E. coli* for 5 days at 20°C, or until populations were one-day old adults. Plates with fungal or bacterial contamination were not included in the assays. Animals were then washed off the plates using S. basal buffer and into microfuge tubes (sterile, but not autoclaved), where they were conditioned to either S. basal buffer (0.1M NaCI, 0.05M K3PO4, pH 6.0), or 2- butanone (Sigma) diluted in S. basal buffer, 1:10,000 (1.23 mM) concentration. Animals were incubated for 80 minutes on a rotating microfuge tube stand. Next, animals were washed two times with S. basal, where the worms were allowed to pellet (about two to three minutes) without spinning them down. Next, the worms were washed with ddH2O to ensure all salts were removed, and then placed onto a 10 cm Petri dish chemotaxis plate. The chemotaxis plate media was made by adding 100 mL ddH2O to 1.6 g of Difco bacto agar, then boiled, and then 500 μl of 1M K3PO4, 100 μl 1M CaCl2 and 100 μl 1M MgSO4 were added, and then the media was pipetted at 10 ml per 10 cm plastic petri dish and let cool to solidify, and then square odor arenas and an origin were drawn (S1 Fig). The chemotaxis plate was made with two point sources – one arena with 1 ul of 200-proof ethanol, and the other arena with 1 μl of 1:1000 butanone in ethanol. Each point source also contained 1 μl of 1M sodium azide (Fisher Scientific) to paralyze worms once they reached the arenas (S1 Fig). Animals were placed at the origin, the liquid was removed with a Kim Wipe, and then animals were allowed to roam for at least two hours. Then, the chemotaxis index or learning index was calculated (Fig 1 legend). For reference, most untrained or buffer-trained wild-type animals have a butanone chemotaxis index from 0.6 to 0.9. If pairwise comparisons between the chemotaxis indices of the buffer-trained and butanone-trained populations of the same genotype were not significantly different from each other, then we deemed them memory defective.

### LTM chemotaxis assay

The assay is performed as written above in the “Chemotaxis assay” section, but for three cycles instead of just one, with recovery periods on food in between. Animals are incubated for 80 minutes in either buffer or butanone diluted in buffer. Animals are washed, and then incubated for 30 minutes in OP50 diluted in S. basal buffer (OD = 10). Animals are washed as before, but then conditioned for another 80 minutes in either buffer or diluted butanone, making this the second odor-treatment cycle. Animals are then washed and incubated with OP50 for another 30 minutes, washed, and then subjected to a third buffer or odor conditioning cycle. Animals are then washed as before, and plated onto chemotaxis plates (this is the 0’ recovery, three-cycle trained worms), or recovered on 5.5 cm NGM plates seeded with OP50 for either 30’, 120’, or 16 hours, where they are then subjected to the chemotaxis assay.

### Cell Identification

All transgenic lines were made in either the *osm-9*(*syb2140*);p*eft-3*::TIR1::mRuby or *osm-9*(*syb2140*) background. The transgenic lines used for cell ID are *osm-9*(*syb2140*); p*eft-3*::TIR1::mRuby[*pocr-4*::ReachR::AID::mKate2 (25 ng/µl);*pofm-1*::GFP (25 ng/µl)], *osm-9*(*syb2140*);p*eft-3*::TIR1::mRuby[*posm-10*::Chrimson::SL2::mCherry (25 ng/µl);*pofm-1*::GFP (25 ng/µl)], *osm-9(syb2140)*; p*eft-3*::TIR1::mRuby[*podr-1::RFP* (10 ng/µl);*pofm-1*::GFP (25 ng/µl)]. *osm-9(syb2140)*;[*psre-1*::HisCl1::SL2::TIR1::mcherry (5 ng/µl);*unc-122*::RFP (25 ng/µl)], and *osm-9(syb2140)*;[p*srh- 142*::HisCl1::SL2::TIR1::mcherry (5 ng/µl);*unc-122::RFP* (25 ng/µl)]. The animals were imaged after the F2 generation was produced. About 10-15 day 1 adult hermaphrodites were put onto 2% agar pads prepared just before imaging. The transgenic animals were paralyzed with 5 mM tetramisole hydrochloride dissolved in M9 buffer [22 mM KH2PO4, 42 mM Na2HPO4, 85.5 mM NaCl, 1 mM MgSO4]. Next, the animals were imaged on a spinning disk confocal microscope under a 60x oil objective at 23°C using NIS Elements. Z-stacks were taken concurrently using 488 nm and 561 nm lasers at 200 ms exposure and 100% power, and were analyzed using FIJI.

The strain used in NeuroPAL cell ID was created by crossing male otIs670 animals with hermaphrodite *osm-9(syb2140)* animals. 10-15 day 1 adult hermaphrodite *osm- 9(syb2140);otls670* animals were placed onto a 2% agar pad and paralyzed using 5 mM tetramisole hydrochloride diluted in M9 buffer. The animals were imaged on a spinning disk widefield confocal under a 40x water objective at 20°C using MicroManager. Z-stacks were taken sequentially; first, the mNeptune2.5 fluorophore was imaged with a 561 nm laser and a 700/75 nm emission filter. Next, the TagRFP-T fluorophore was excited using the 561 nm laser. Third, CyOFP was imaged using a 488 nm laser with a 605/70 nm emission filter. Fourth, the GFP channel Z-stacks were taken using the 488 nm laser with a 525/50 nm emission filter, and finally the mTagBFT2 was excited using a 405 nm laser. All of the channels were imaged using 50 ms exposure. All 6 channels were imaged using 20% power. The acquired images were analyzed using FIJI, in accordance with the Hobert Lab’s guide “Using NeuroPAL for Neural ID” from Columbia University [58].

### Imaging OSM-9 Expression Pattern before and after K-NAA and in the ZIF-1 degradation system

Animals were staged as described in the short-term chemotaxis assay methods above. Naive N2 and *osm-9(syb2140);TIR1* animals were imaged using NemaGel Beadless (NGEL-200BL). NemaGel was dotted onto a slide and flattened with a coverslip on top; then, the coverslip was quickly removed to create a smooth surface of gel. Next, 5 mM tetramisole at 37°C was washed three times over the surface of the gel. Finally, the animals were washed off an NGM plate with S. basal and pipetted onto the gel, and a coverslip was placed on top. K-NAA (Fisher Sci. Catalog No. N000625G) was dissolved in S-Basal buffer to produce a 100 mM stock and then diluted to 10 and 25 mM. Animals were incubated in either 0 mM, 10 mM, or 25 mM K-NAA buffer while rotating for 2 hours and were then imaged using the same protocol as the naive animals above. Images were analyzed using ImageJ by creating a sum slices Z-projection of either the cilia or the cell bodies (60 slices total for cilia and 35 slices total for cell bodies), using the freehand selection tool to circle the cilia or the cell bodies, and measuring the mean gray value of the selected area. Next, a circle from either directly behind the cilia or directly next to the cell bodies was measured for background mean gray value, and this value was subtracted from the mean gray value of the cilia or cell bodies.

### Sleep analysis

To make the WorMotel (<u>Churgin *et al.*, 2017</u>), we used a PDMS chip with 48 total wells made according to the paper or the online resource (http://fangyenlab.seas.upenn.edu/links.html). Next, we made 100mL of NGM by adding together 1.8g low melting-point agarose, 0.3g NaCl, 0.25g bacto peptone, and 1µL tween 20 (to keep a flat agar surface). We boiled the media in the microwave, cooled it down to ∼50-58°C, then added 100µL CaCl2, 100 µL cholesterol dissolved in EtOH, 100 µL MgSO4, and 2.5 mL K3PO4. We next added 17 µL/well of chip and let it cool. The solidified WorMotel was placed in a transparent Petri dish with 10mg of gel soil (Soil Moist granules) soaked in 100 mL of water to prevent cracking of wells. Four clay balls were used to prop the lid open uniformly for 1-8 hours or as long as the worms were assayed. Food from NGM/OP50 plate was scooped and smeared on top of the wells to prevent the worms from being starved. Teledyne Dalsa PT-21-04M30 Camera Link Camera (Dalsa Proprietary Sensor 2352x1728 Resolution Area Scan Camera) attached with a Linos Rodagon Modular-Focus lens (f=60mm) was used to image the entire WorMotel. To obtain a focused working distance, four metal posts with a plastic stage was built. A T175 tissue culture flask filled entirely with water was added to make a cooling chamber as well as a light diffuser (water diffracts light). We used the Multiple-Worm Tracker (MWT 1.3.0r1041) made by Rex Kerr to automate image capture and record worm movement every 3 seconds for the entire duration of the experiment.

Irfanview (developed by Irfan Škiljan) was used to re-index the images for sequence verification before quantifying the movements in MATLAB. Once indexed, the images were batch processed in MATLAB for thresholding each worm uniformly to quantify quiescence using a graphic user interface (GUI) created by MC and CFY and available at https://github.com/LEtoileLab/Sleep_2022.git. After thresholding, the animals were quantified for quiescence and activity using another MATLAB code available at https://github.com/LEtoileLab/Sleep_2022.git

### Statistical Analysis

All data included in the same graph were subjected to the Shapiro-Wilk normality test. If all of the datasets were normally distributed, then one-way ANOVA was performed, followed by Bonferroni correction for pairwise comparisons. If any datasets were non-normally distributed, then the Kruskal-Wallis test was performed. If p>0.05, then no further analysis was performed. If p<0.05, then the test was followed up by either the Mann-Whitney u-test for non-parametric data pairwise comparisons or the Student’s unpaired t-test for parametric data pairwise comparisons. All p-values included in the same graph were then adjusted using the Hochberg test to remove any type I statistical error, which prevents incorrect rejection of the null hypothesis. ***p<0.001, **p<0.01, *p<0.05, NS indicates p>0.05. All graphs show S.E.M. Graphpad Prism and R studio were used for all of the statistical tests.

Each data point (represented by grey dots) on the graphs indicates one population of 400>N>50, run on independent days.

## Acknowledgements

We would like to thank all past and present members of the L’Etoile lab. We thank Sarah Hall for her gift of the SH272 *osm-9* whole-gene deletion strain (used in a previous version of this ms), Piali Sengupta for her gift of the *nphp-4* promoter and Cori Bargmann for her gift of the *osm-9::GFP5* construct. We would also like to thank Joe Hill for the artwork used in S1 and S2 Figs. We thank the Caenorhabditis Genetics Center (CGC), supported by the NIH Office of Research Infrastructure Programs (P40 OD010440), for the strains they provided for this study. We are grateful to the team behind wormbase.org (NIH grant #U24 HG002223), SMART (Schultz et al., 1998, EMBL), and Nikhil Bhatla for the wormweb.org exon-intron graphic maker used for Fig 7A. For funding support we would like to thank: KB: F31 DC014921 and R01 supplement NS87544; MFS: F31DC019872; RLD: F31NS115572; SK: R35GM124735; NDL: R01 NS087544 and R01DC005991.

## Supporting information

**S1 Fig.**
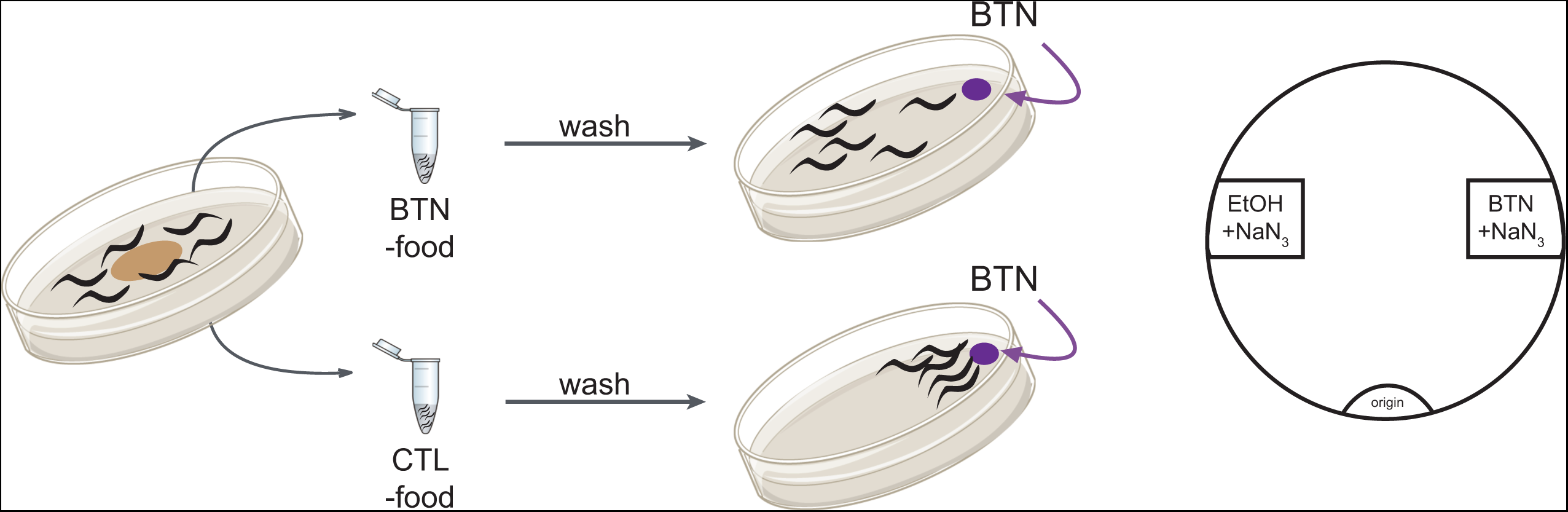
Schematic of one-cycle odor-conditioning. Animals are conditioned to butanone as previously published [51]. In brief, age-synchronized, one-day old adult animals are washed with buffer from NGM plates containing OP50 food to tubes with either buffer or butanone, both containing no food, for 80 minutes. After the 80-minute incubation, animals are washed and then plated onto chemotaxis assay plates. The chemotaxis index (See Fig 1 legend) is then calculated after at least two hours of the animals roaming.

**S2 Fig.**
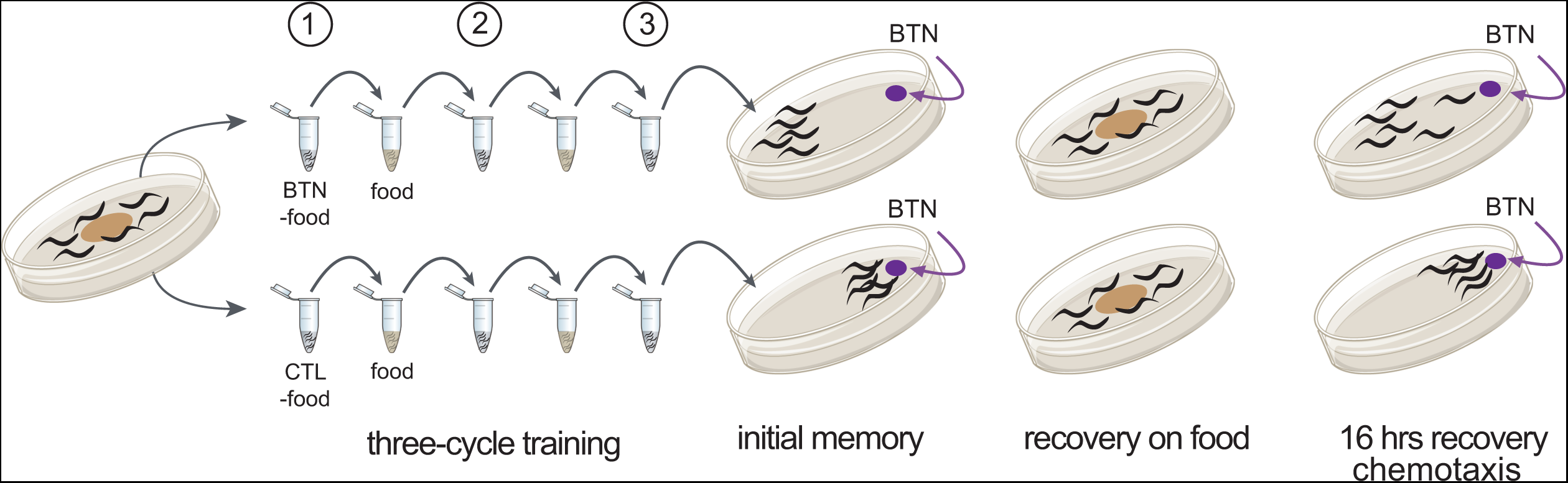
Spaced-training paradigm. Schematic of associative olfactory memory assay to induce long-term odor memory. Age-synchronized, one-day old adult animals are washed from plates with food into tubes and are conditioned to either buffer (CTL) or butanone (BTN) for 80 minutes per cycle in the absence of food for three cycles total, with a 30-minute recovery on food (OP50, OD = 10) in between cycles. After spaced-training, animals are either tested in a butanone chemotaxis assay or are recovered on OP50-seeded NGM plates and then tested.

**S3 Fig.**
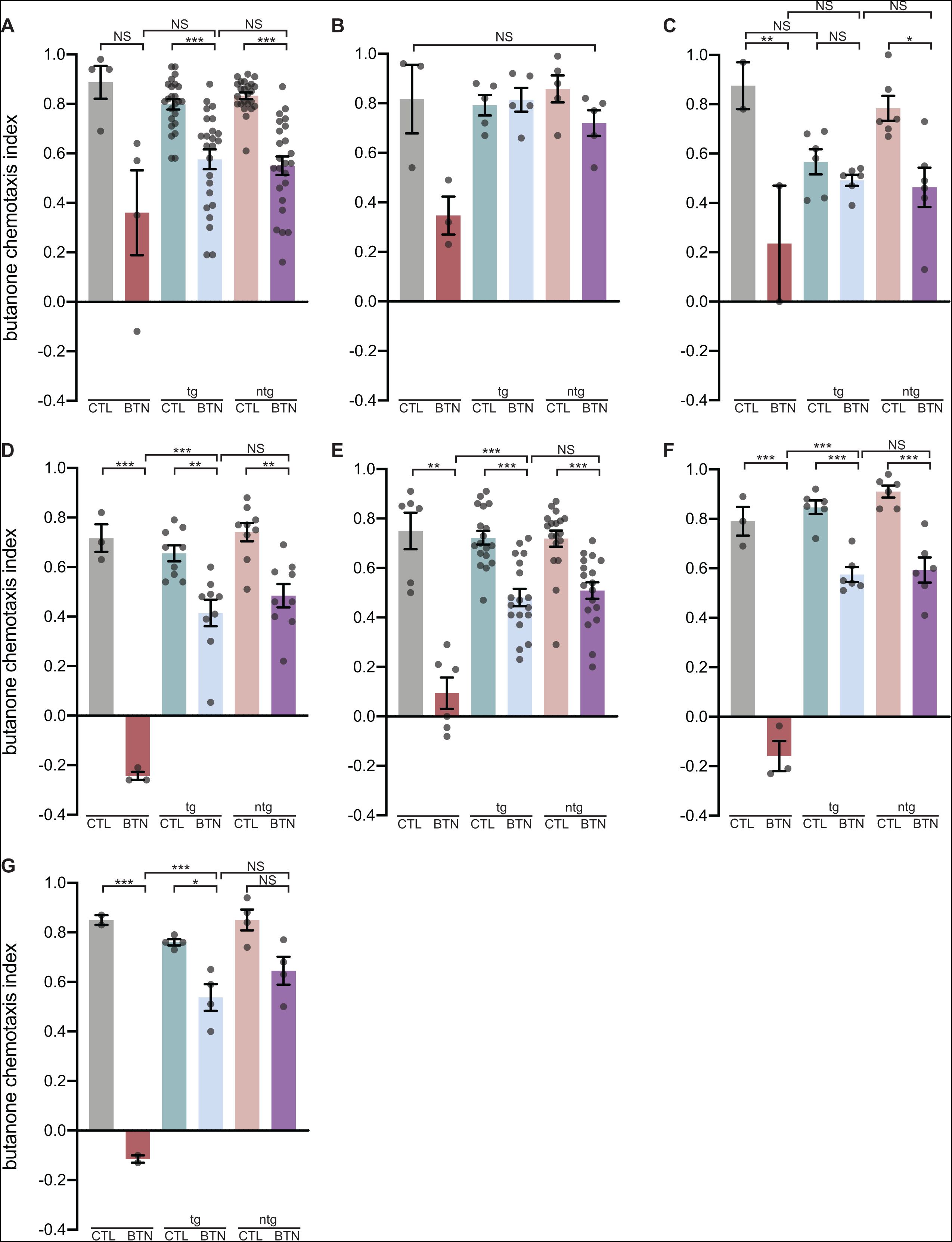
Various *osm-9* expression transgenes are not sufficient to rescue *osm- 9(ky10)* one-cycle butanone conditioning defects. (A) Chemotaxis indices of wild type versus *osm-9(ky10),* with or without the transgene *KBC8/pceh-36::osm- 9SS::GFP::osm-9.* The *pceh-36* is a promoter driving expression in both of the AWC olfactory sensory neurons. The *osm-9SS* (signal sequence) cDNA was 5’ of the GFP and the rest of the cDNA was 3’ of the GFP. Data shown from six combined transgenic lines. The Kruskal-Wallis test was performed. Transgenic is tg and non-transgenic is ntg. The u-test was used for CTL vs BTN and ntg CTL vs ntg BTN. The t-test was performed for tg CTL vs tg BTN, BTN vs tg BTN, and tg BTN vs ntg BTN. (B) Chemotaxis indices of wild type versus *osm-9(ky10),* with or without the transgene *pJK21/podr-3::GFP::osm-9.* The *podr-3* promoter drives expression in all three of the pairs of olfactory sensory neurons, including AWA, AWB, and AWC. The *podr-3* used was the short promoter, 2653 bp in length. The *osm-9* used in the same partial cDNA as in panel A. Data shown from two combined transgenic lines. The Kruskal-Wallis test was used and p>0.05. (C) Chemotaxis indices of wild type versus *osm-9(ky10),* with or without the transgene *KBC24/podr-3::GFP::osm-9.* The *podr-3* promoter drives expression in all three of the pairs of olfactory sensory neurons, including AWA, AWB, and AWC. The *podr-3* used was the long promoter, 2686 bp in length. The *osm-9* used in the same partial cDNA as in panel A. Data shown from three combined transgenic lines. One-way ANOVA was performed, followed by Bonferroni correction for CTL vs BTN, tg CTL vs tg BTN, ntg CTL vs ntg BTN, BTN vs tg BTN, tg BTN vs ntg BTN, CTL vs tg CTL, and tg CTL vs ntg CTL. (D) Chemotaxis indices of wild type versus *osm-9(ky10),* with or without the transgene *osm-9::GFP5* [37], which includes the *osm-9* partial cDNA (2497 bp, missing the last 317 bp) driven by the upstream ∼1.6 kb promoter and downstream ∼3.2 kb promoter, and GFP, cloned into the pBS parent vector. Data shown from three combined transgenic lines. One-way ANOVA was performed, followed by Bonferroni correction for CTL vs BTN, tg CTL vs tg BTN, ntg CTL vs ntg BTN, BTN vs tg BTN, and tg BTN vs ntg BTN. (E) Chemotaxis indices of wild type versus *osm-9(ky10),* with or without the transgene *KBC42/posm-9::osm-9::UTR*, where *posm-9* is the upstream ∼1.6 kb promoter, *osm-9* is the full-length cDNA (2814 bp), and *UTR* is the ∼3.2 kb downstream promoter. Data shown from three combined transgenic lines. The Kruskal-Wallis test was performed. The u-test was used for CTL vs BTN and ntg CTL vs ntg BTN. The t-test was performed for tg CTL vs tg BTN, BTN vs tg BTN, and BTN tg vs ntg BTN. (F) Chemotaxis indices of wild type versus *osm-9(ky10),* with or without the transgene WRM065bE12, a ∼32 kb fosmid containing the entire *osm-9* gene its upstream and downstream UTR regions. Data shown from two combined transgenic lines. One-way ANOVA was performed, followed by Bonferroni correction for CTL vs BTN, tg CTL vs tg BTN, ntg CTL vs ntg BTN, BTN vs tg BTN, and tg BTN vs ntg BTN. (G) Chemotaxis indices of wild type versus *osm-9(ky10),* with or without the transgene *KBC38/pnphp-4::osm-9::GFP.* The *pnphp-4* promoter drives expression in all of the ciliated neurons (∼60) in the adult worm. The *osm-9* transgene was the full-length cDNA. Data shown from two combined transgenic lines. One-way ANOVA was performed, followed by Bonferroni correction for CTL vs BTN, tg CTL vs tg BTN, ntg CTL vs ntg BTN, BTN vs tg BTN, and tg BTN vs ntg BTN.

**S4 Fig.**
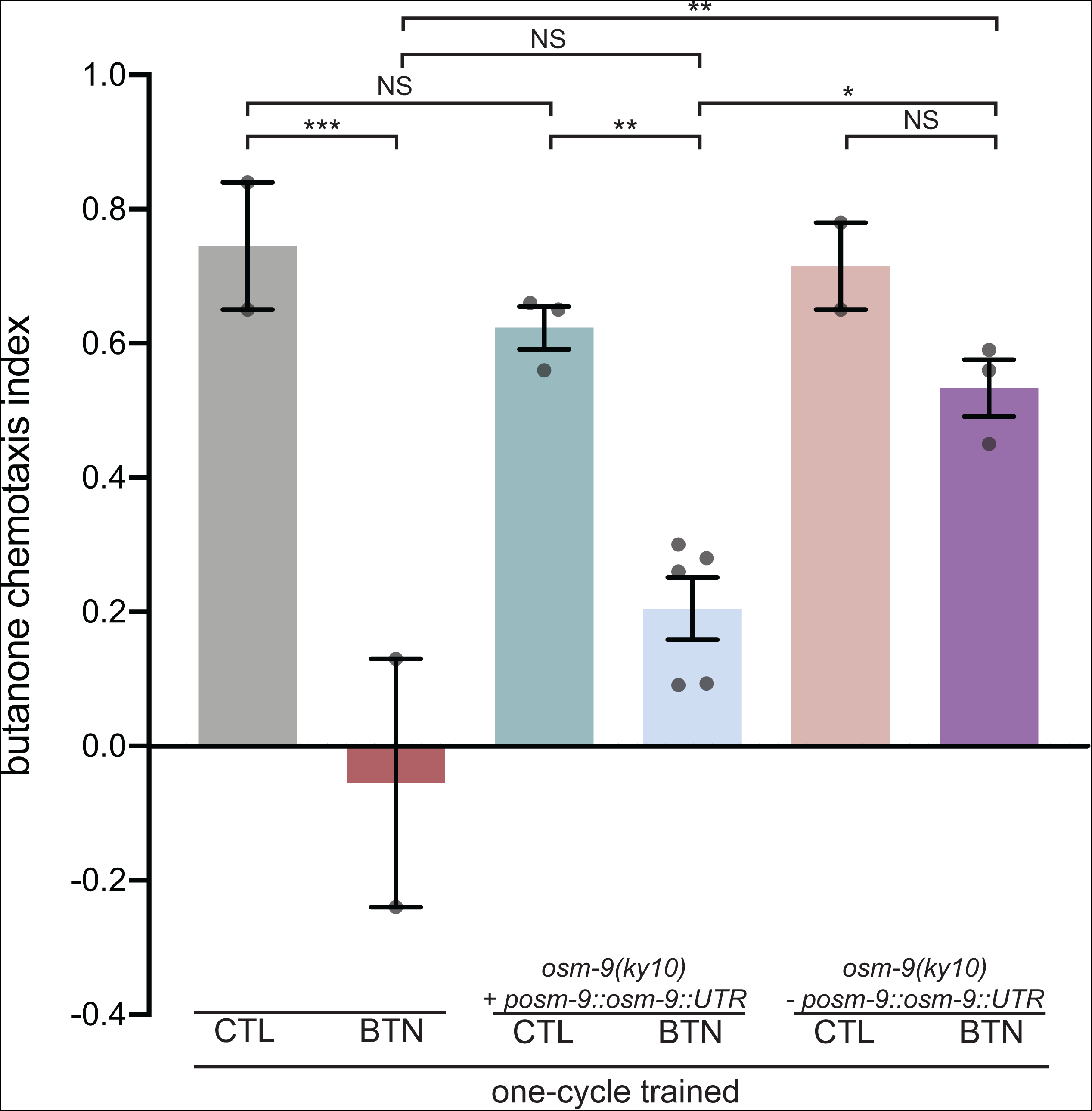
Expressing *osm-9* gDNA with its upstream and downstream endogenous sequences in *ky10* rescues short-term memory. Chemotaxis indices of one-cycle trained animals. *KBC56/posm-9::osm-9::UTR* is the rescue transgene, where *posm-9* is the 1.6 kb upstream promoter (Fig 7A), *osm-9* is the full-length gDNA (6342 bp), and UTR is the 3.2 kb downstream sequences (possibly enhancer) (Fig 7A). Teal and blue bars are *osm-9(ky10)* animals expressing *posm-9::osm-9::UTR* and pink and purple bars are the *osm-9(ky10)* non-transgenic siblings. Uneven number of grey dots because only datasets with N>50 were counted in the graph and analysis. One-way ANOVA was performed, followed by Bonferroni correction for CTL vs BTN, *osm-9(ky10) + posm- 9::osm-9::UTR* CTL vs *osm-9(ky10) + posm-9::osm-9::UTR* BTN, *osm-9(ky10) - posm- 9::osm-9::UTR* CTL vs *osm-9(ky10) - posm-9::osm-9::UTR* BTN, BTN vs *osm-9(ky10) + posm-9::osm-9::UTR* BTN, *osm-9(ky10) + posm-9::osm-9::UTR* BTN vs *osm-9(ky10) - posm-9::osm-9::UTR* BTN, CTL vs *osm-9(ky10) + posm-9::osm-9::UTR* CTL, and BTN vs *osm-9(ky10) - posm-9::osm-9::UTR* BTN.

## Notes

### Competing Interest Statement

The authors have declared no competing interest.

### Summary of Updates

We have added new data in figure 3 that shows ism-9 mutants sleep like wild types. New data showing that loss of the ankyrin domains does not affect memory. New data showing that chronic loss of OSM-9 in some ciliated neurons blocks memory consolidation. Four new authors helped with these data and were added.

